# A Population Genomic Assessment of Three Decades of Evolution in a Natural *Drosophila* Population

**DOI:** 10.1101/2021.12.23.474033

**Authors:** Jeremy D. Lange, Héloïse Bastide, Justin B. Lack, John E. Pool

## Abstract

Population genetics seeks to illuminate the forces shaping genetic variation, often based on a single snapshot of genomic variation. However, utilizing multiple sampling times to study changes in allele frequencies can help clarify the relative roles of neutral and non-neutral forces on short time scales. This study compares whole-genome sequence variation of recently collected natural population samples of *Drosophila melanogaster* against a collection made approximately 35 years prior from the same locality – encompassing roughly 500 generations of evolution. The allele frequency changes between these time points would suggest a relatively small local effective population size on the order of 10,000, significantly smaller than the global effective population size of the species. Some loci display stronger allele frequency changes than would be expected anywhere in the genome under neutrality – most notably the tandem paralogs *Cyp6a17* and *Cyp6a23*, which are impacted by structural variation associated with resistance to pyrethroid insecticides. We find a genome-wide excess of outliers for high genetic differentiation between old and new samples, but a larger number of adaptation targets may have affected SNP-level differentiation versus window differentiation. We also find evidence for strengthening latitudinal allele frequency clines: northern-associated alleles have increased in frequency by an average of nearly 2.5% at SNPs previously identified as clinal outliers, but no such pattern is observed at random SNPs. This project underscores the scientific potential of using multiple sampling time points to investigate how evolution operates in natural populations, by quantifying how genetic variation has changed over ecologically relevant timescales.

## Introduction

A central goal in population genetics is to understand the relative contributions of neutral and non-neutral forces on genetic variation. Typical genome-wide analyses of genetic variation examine these forces across a wide interval of evolutionary time by examining a single snapshot of genetic variation, reflecting events from roughly the last 4*N*_*e*_ generations (where *N*_*e*_ is the effective population). Adaptation, however, can occur over much shorter ecological timescales in natural populations (Daborn *et al*. 2001, Colosimo *et al*. 2005, Hoekstra *et al*. 2006, Campbell-Staton *et al*. 2017, Pélissié *et al*. 2018). While researchers have long sought to understand short-term adaptation, decreasing sequencing costs in recent years have sparked a renewed interest in short-term adaptation studies.

Utilizing multiple sampling times to study temporal changes in allele frequencies can clarify the relative roles of neutral and non-neutral forces on very short time scales. This technique helps minimize neutral genetic differences when comparing genomic variation between time points. For example, evolve and resequence (E&R) studies use a multiple sampling time technique by evolving laboratory populations in controlled environments and observing changes in genetic variation over perhaps dozens of generations. E&R studies have been conducted on multiple species including *E. coli* (Barrick *et al*. 2009), influenza (Foll *et al*. 2014), *S. cerevisiae* (Parts *et al*. 2011), and *D. melanogaster* (Turner *et al*. 2011) to uncover important dynamics and targets of natural selection. One disadvantage of E&R studies is that they may not always reveal how selection works in nature because they are typically derived from laboratory populations. Laboratory populations generally contain only a subset of natural genetic diversity, and are kept in environments free of natural enemies, and are typically exposed to only specific controlled environmental stresses (potentially minimizing the pleiotropic consequences of laboratory adaptation).

There have now been a multitude of temporal population genomic studies in humans (Burger *et al*. 2007, Mathieson *et al*. 2015, Hofmanová *et al*. 2016) and other mammals (Noonan *et al*. 2005, Lindqvist *et al*. 2010, Castañeda-Rico *et al*. 2020). Short-lived organisms such as *Drosophila melanogaster*, which goes through hundreds of generations within a single human generation (about 15 generations per year; Turelli and Hoffmann 1995; Pool 2015), allow the study of substantially more evolution over shorter time scales. A study by Bergland *et al*. (2014) found evidence for dozens of genomic loci showing seasonal allele frequency changes in *D. melanogaster* that could not be explained by drift alone. Subsequent studies have expanded on those findings by leveraging seasonal population genomic data across multiple years from dozens of locations (Kapun *et al*. 2020; Machado *et al*. 2021). Researchers have also sequenced museum specimens of the North American honeybee to study genomic changes in response to the introduction of a parasitic mite across roughly 50 generations (Mikheyev *et al*. 2015). Feder *et al*. (2016) used multiple sampling time points in humans to examine the roles of hard and soft selective sweeps in HIV over short time scales. More recently, Chen *et al*. (2019) examined a natural pedigreed population of 3984 Florida scrub jays over a period of 24 years (∼5 generations). The authors discovered several SNPs under directional selection during this brief interval, suggesting the importance of rapid adaptation in natural populations.

In *D. melanogaster*, a few studies have examined long-term frequency changes in specific polymorphisms (*e*.*g*. Umina *et al*. 2005; Kapun *et al*. 2016). However, none have examined changes in genome-wide genetic variation across multiple decades in this population genetics model. The research described here compares genomic variation of a recently collected sample of *D. melanogaster* to samples from the same locality collected approximately 35 years ago (∼500 generations ago). The collection of flies from 35 years ago has been maintained in a lab as 65 independent isofemale strains since their collection. Because it is prohibitively unlikely that the same mutation has occurred in many individual strains, and because selection on the limited variation present at sampling is inefficient in these small lab cultures, the genetic variation we observe across these 65 strains should represent an accurate representation of genetic diversity in this population 35 years ago. This study gives us an unusually direct way of quantifying change in genomic variation across decades, allowing us to ask important questions about how evolution works in nature in a populous insect species. Because we can focus on a much narrower time scale than typical population genomic analyses, we can begin to examine how much selection has occurred over the last ∼35 years. We also investigate regions of the genome that may have undergone very recent selection, which could inform more precise investigations into genes associated with, for example, insecticide resistance or climate adaptation.

## Results

### Genome Sequencing and Quality Control

Whole genome sequences were extracted from 65 wild-derived isofemale strains of *Drosophila melanogaster* originally collected from Providence, Rhode Island (USA) between 1975 and 1983. Mean sequencing depth per individual genome ranged from 5.3X to 72X, with an average of 24X (Table S1). We returned to Providence in Fall 2014 and Spring 2015 to collect two distinct seasonal samples. Pooled sequencing was used to maximize the number of flies that could be included and hence the precision of allele frequency estimates. From each wild-collected female, we included one F1 daughter in the pool if morphological examination of that same female’s sons confirmed species identity. 247 flies divided into 6 pools were sequenced in the fall sample (with a total mean depth of 384X) and 408 flies divided into 18 pools were sampled in the spring sample (with a total mean depth of 194X; Table S2).

We applied two quality control approaches to confirm the expected genetic composition of the old isofemale lines. First, we performed a pairwise analysis of identity-by-descent (IBD). One pair of genomes was found to be essentially identical, apparently reflecting an accidental stock duplication, and one member of this pair was therefore excluded from subsequent analysis. Other minor instances of apparent relatedness were also noted and masked (Table S3). As a further check for contamination, we checked IBD was found between Providence strain genomes and the common laboratory genetic background represented by the *D. melanogaster* reference strain. No large-scale IBD was found outside regions of known recurrent IBD; the largest blocks spanned the chromosome 3 centromere, which often shows prominent IBD between independent strains from the same region (Lack *et al*. 2015).

We also performed Principle Components Analysis (PCA) to test for any genetic outliers among the old strains and confirm their expected relationships to other populations. All of the old Providence genomes clustered with other northeast US genomes (New York), and between those from North Carolina and those from France (Figure 1), as anticipated based on a known latitudinal cline of African versus European ancestry (Kao *et al*. 2015; Bergland *et al*. 2016). We further investigated whether the old Providence population sample showed any unusual pattern of variance among individuals with regard to the PCA results. We initially focused on the within-population standard deviations of principle components PC1 and PC2, because their weightings corresponded to 7.3% and 1.6% of the variation among individuals, respectively, whereas PC3 through PC32 only explained 0.4 – 0.6% each (Table S4). We found that the Providence sample had SD(PC1) and SD(PC2) values intermediate to the other analyzed population samples (Figure S1). Focusing on the other two larger samples, the European FR sample from France and the North Carolina RAL population with somewhat greater African ancestry than Providence, we find that Providence had significantly lower SD(PC1) than France but significantly higher SD(PC1) than RAL; whereas for SD(PC2) we find that Providence had modestly but significantly lower SD(PC2) than either FR or RAL (bootstrap-resampled *P* < 0.0001 in each case). Similar results were obtained when assessing the proportion of PC outliers for Providence compared with FR or RAL (the samples with adequate sizes for this analysis), with outliers defined as the proportion of individuals more than two population SDs from the population mean. For both PC1 and PC2, Providence’s outlier proportion was intermediate to FR and RAL (Figure S1), and none of the differences were statistically significant (z test *P* > 0.5 in each case). These results suggest that the old Providence genomes show levels of genetic heterogeneity consistent with those expected for a geographically uniform North American population sample of *D. melanogaster*.

**Figure 1.**
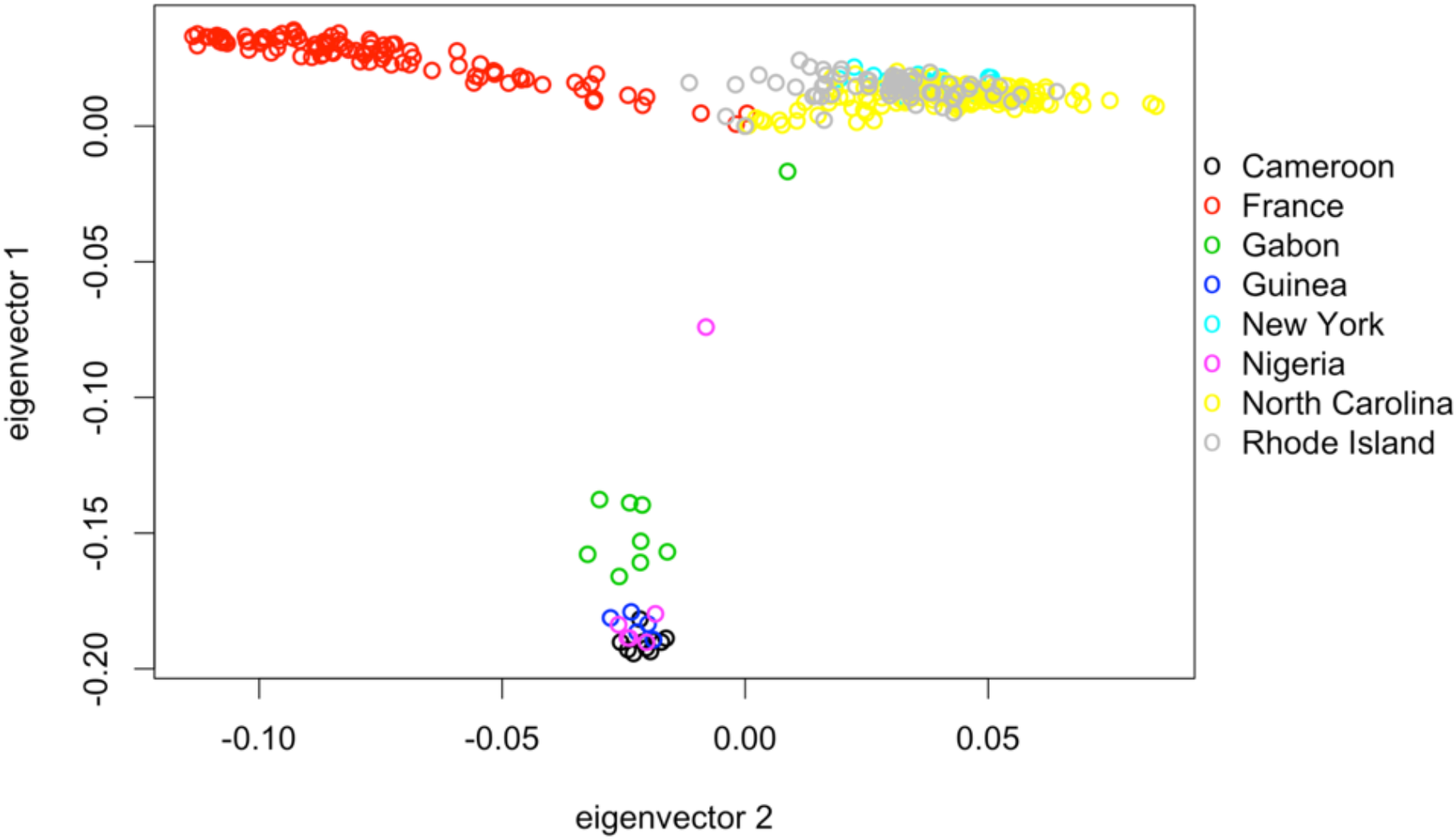
Principle Components Analysis reflects expected relationships between old Providence genomes and other sequenced genomes, with no obvious outlier individuals. X chromosome SNP data from 7 additional North American, European, and African populations were used in this analysis.

### Inversion Analysis

Because chromosomal inversions suppress recombination in a heterozygous state (Sturtevant and Beadle 1936), they have long been believed to play important roles in evolutionary processes. Indeed, research has established that inversions can play roles in natural selection (Kirkpatrick and Barton 2006; Kapun *et al*. 2016) and reproductive isolation (Noor *et al*. 2001). *D. melanogaster* inversions are also known to affect genetic diversity over broad chromosomal scales (Corbett-Detig *et al*. 2012; Pool *et al*. 2012). We therefore wanted to measure common inversion frequencies in both the old and new samples.

As a reference point, we note that Mettler *et al*. (1977) assayed inversion frequencies from wild-collected flies including from Portland, Maine and Niagara Falls, New York. In these two populations, they found the fraction of autosomal chromosome arms carrying an inversion to be 3.98% and 0.0125%, respectively. In contrast, our old Providence strains had 12.1% inverted autosomal arms. While geographic variation in inversion frequencies is possible, another possibility for our lab-maintained strains is associative overdominance – that because inversion polymorphism can keep large chromosomal regions heterozygous, it may be selectively maintained in small lab cultures by buffering against recessive deleterious variants, which are common on randomly sampled chromosomes from wild populations (Greenberg and Crow 1964). By comparison, our new fall and spring samples had 9.06% and 8.72% inverted autosomal arms, respectively. Although these frequencies are slightly lower than our old sample, we can not rule out the possibility that inversions have actually increased in frequency through time, in light of the Mettler *et al*. (1977) data and the associative overdominance hypothesis described above.

Seasonal change in *Drosophila* inversion frequencies is a long-standing topic of study (*e*.*g*. Dobzhansky 1943). We therefore asked whether inversion frequencies varied seasonally between our fall and spring samples (Table S5). Here, we simulated sampling of individuals and reads based on empirical SNP frequencies and empirical depth of coverage at inversion-associated SNPs and asked how often we observed inversion frequency differences between seasons as large as what we observed in the real data (see Materials and Methods). We found that sampling variation could not explain seasonal inversion frequency differences for three inversions: *In(2R)NS, In(3R)Mo*, and *In(3R)P*. The observed frequency of *In(2R)NS* decreased from 0.068 in the fall to 0.029 in the spring, a shift that our resampling simulation indicated was unlikely to be due to sampling variance (p=0.0033 based on simulations; see Materials and Methods). In contrast, *In(3R)P* rose in frequency from 0.0394 in fall to 0.0887 in spring, with an associated P value of 0.0016. A second inversion on the same chromosome arm, *In(3R)Mo*, also increased, from 0.055 in fall to 0.0895 in spring, for an associated P value of 0.0301. The frequency differences we observed in other common inversions (*In(2L)t, In(3L)P, In(3R)C*, and *In(3R)K*) could all be explained by sampling variance. Our results are mostly in line with Kapun *et al*. (2016), who also described significant seasonal differences in *In(2R)NS* and *In(3R)P*, but not *In(3R)Mo*, from eastern US populations. Whereas, Machado *et al*. (2021) analyzed the association between seasonally-variable SNPs and inversion breakpoint regions, finding significant associations for *In(2L)t* and for a set of three inversions on arm 3R, but not for *In(2R)NS*.

Although we appear to have observed real frequency changes at these three inversions, it is unclear why. One hypothesis is that there was drift caused by seasonally reduced winter population size. Under such a model, one could expect inversion frequency differences between sampling locations in the spring sample, since local population sizes should be lower leading up to the spring samples than for the fall samples. We thus asked whether inversions differed in frequency between our five sampling locations (which are all within 1.7 km of each other) within each season. Here, we employed a similar simulation-based sampling strategy as for the seasonal inversion analysis above. Instead of asking how often we observed seasonal differences as large as we observed empirically, we asked how often we observed inversion frequency differences between sampling locations as large as what we observed in the real data (see Materials and Methods). We did not find evidence of inversion frequency differences between sampling locations in the fall sample; all differences could be explained by sampling variance. This was not the case for the spring data (Figure 2B). All three inversions where we found significant seasonal differences (*In(2R)NS, In(3R)Mo*, and *In(3R)P*), also showed significant frequency differences between sampling locations. Additionally, *In(3R)C* also showed significant differences between sampling locations, although its seasonal difference could be explained by sampling variation (p=0.703). Our observed seasonal and spatial differences in inversion frequencies could potentially be explained by genetic drift, due to low population sizes coming out of a cold winter season. This explanation is consistent with the higher genome-wide genetic differentiation observed between spring sample locations (mean *F*_*ST*_ = 0.046) than between fall samples (mean *F*_*ST*_ = 0.009; Table S6). However, it remains plausible that some inversions may have seasonally-variable fitness consequences, in line with the parallel findings from our study and Kapun *et al*. (2016) for *In(2R)NS* and *In(3R)P*, and between our study and Machado *et al*. (2021) for arm 3R inversions, although the specific patterns observed may depend on geography and other factors.

**Figure 2.**
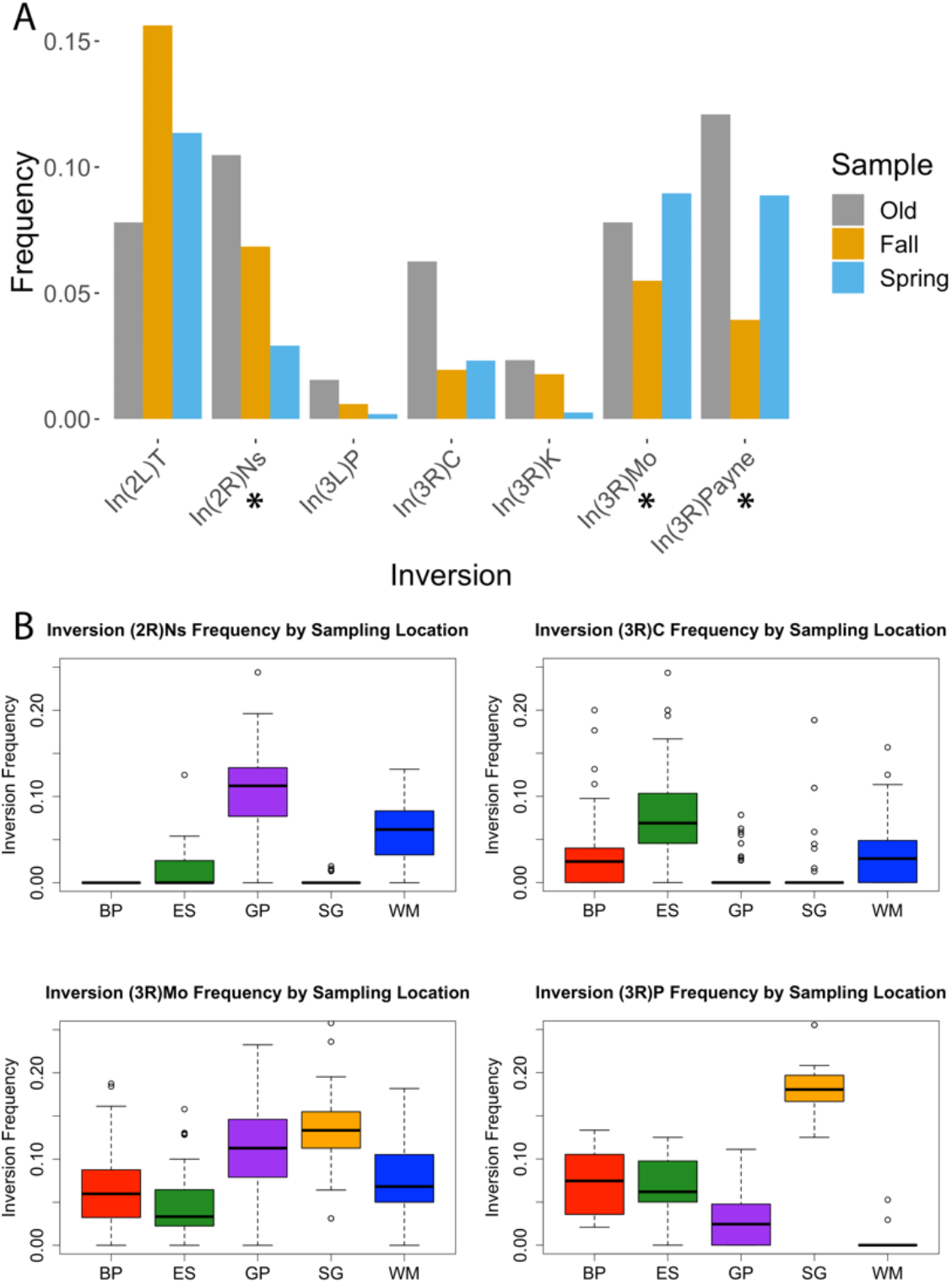
Inversion frequencies have shifted across time and space. (A) Inversion frequencies at 8 well-studied chromosomal inversions. The asterisks underneath *In(2R)Ns, In(3R)Mo*, and *In(3R)P* indicate statistically significant frequencies between seasons. (B) Four inversions showed significant frequency differences between sampling locations (x-axis) in the spring, as detailed in Table S2. No inversions displayed significant frequency differences between locations in the fall samples.

### Genome-wide Ancestry may have Slightly Shifted Toward European Alleles

North American populations of *D. melanogaster* were founded roughly 150 years ago (Keller 2007) and involve an admixture event between a majority European-like gene pool and a minority African-like gene pool (Caracristi and Schlötterer 2003; Duchen et al. 2013; Pool 2015). This admixture may have resulted from secondary contact between African migrants introduced to the Caribbean (or neighboring regions) and European migrants potentially introduced to the northeast US (Keller 2007), ultimately generating an ancestry cline along the east coast of North America (Kao *et al*. 2015; Bergland *et al*. 2016). Such admixture can have evolutionary consequences by, for example, introducing novel adaptive variants (*e*.*g*. Racimo *et al*. 2015) or generating epistatic interactions between alleles of different ancestry (*e*.*g*. Pool 2015). It is therefore important to estimate proportions of African and European ancestry in this population at the two time points. A significant change in genome-wide ancestry over the 35-year time period could signal a shift in the east coast ancestry cline, potentially due to demographic effects such as asymmetric migration.

Overall, we estimated a genome-wide median of 17.26% African ancestry across tested SNPs in the old genomes. We found little variability in ancestry proportion from one genome to the next in the old samples, with a standard deviation of 2.30% on chromosome 2, 3.98% on chromosome 3, and 4.04% on chromosome X. This variability from genome to genome is similar to previous estimates from inversion-free chromosomes from the *Drosophila* Genetic Reference Panel (DGRP) population that originates from North Carolina (Pool 2015), if we account for the slightly higher mean African ancestry among DGRP genomes.

We estimated somewhat lower genome-wide levels of African ancestry in the newer samples, with a median of 13.43% in the 18 seasonal pools. The ancestry estimate in the old samples may be skewed if inversion frequencies shifted within the isofemale lines since the original collection, as suggested above. We observed a decrease in frequency in 6 of 7 tested inversions. If the higher inversion frequencies in our old strains can be attributed to associative overdominance (as suggested above), instead of a true decrease of inversion frequencies between time points, then the African ancestry we measure in the old samples may also be overestimated because of the association between African ancestry and inversions (Corbett-Detig & Hartl 2012; Pool 2015). To help control for differences in ancestry between time points, we weighted individual samples in the old samples to match inversion frequencies in the pooled sequences.

Using this approach, we observed a slightly lower African ancestry in the old samples (15.94%, Figure 3). Thus, after controlling for inversion frequencies, we still estimate a decline in genome-wide African ancestry between time points: roughly 2.5% in absolute terms, or in relative terms, a loss of 15.7% of the African ancestry that was present in the old sample. Explanations for this genomic ancestry shift could include directional selection (*e*.*g*. an advantage of European alleles in this temperate environment), epistatic selection (against incompatible African variants), and population history (*e*.*g*. asymmetric migration).

**Figure 3.**
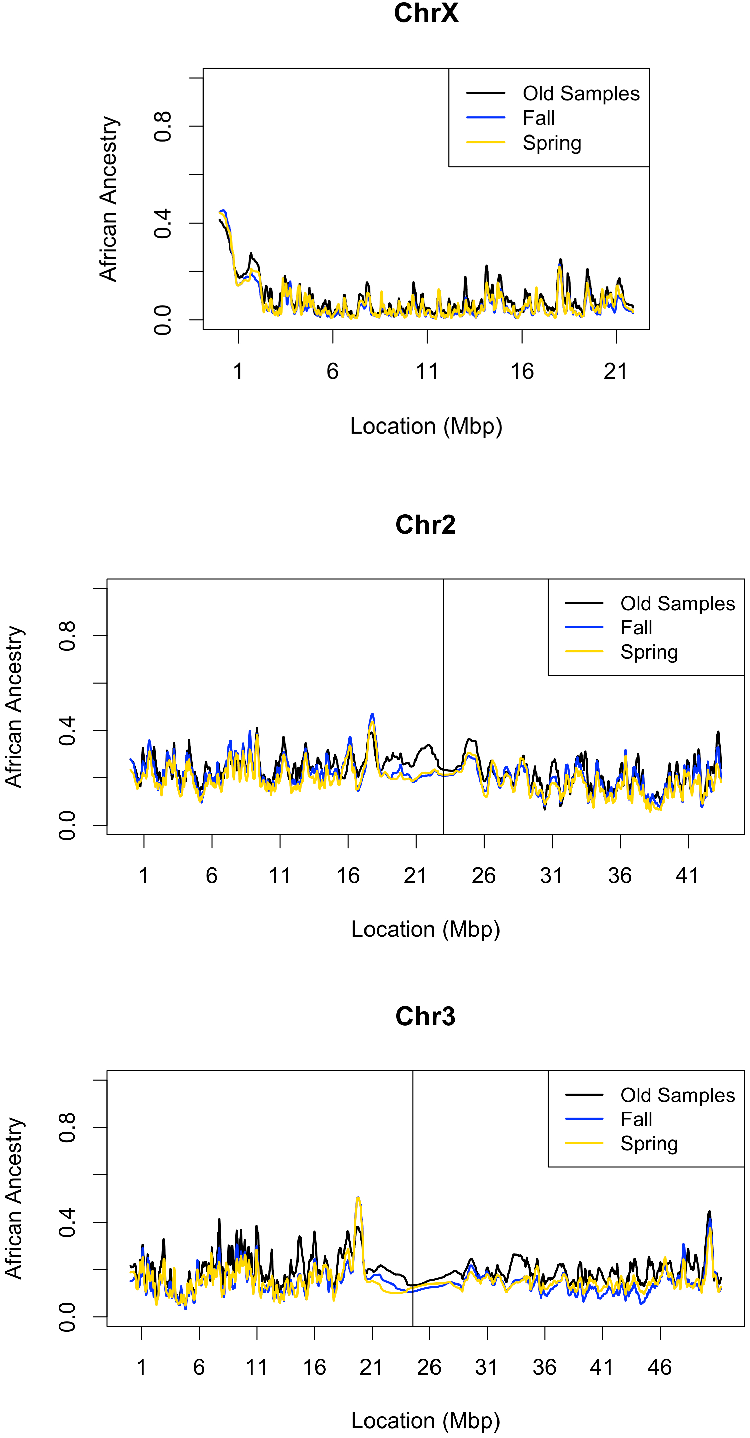
Population ancestry appears to have not shifted appreciably between time points. African versus European ancestry was estimated along chromosomes using the hidden Markov model of Corbett-Detig & Nielsen (2017). African ancestry proportion was averaged across inversion-free strains in the original samples (black line), averaged across 6 pools in the fall sample (blue), and 12 pools in the spring sample (yellow). Because inversions can affect ancestry, and we observed shifts in inversions frequencies between time points, we weighted ancestry in the old samples to match inversion frequencies between the time points. This weighted ancestry along the genome is shown here.

We can complement this genome-wide examination of the ancestry cline by looking for SNP frequency changes at highly clinal outlier SNPs that may reflect the action of spatially-varying selection. Using the data set provided from Machado *et al*. (2016), we examined the temporal dynamics of all SNPs with a clinal P value below 0.001. By cross-referencing these SNPs with the data set provided in Bergland *et al*. (2014), we were able to identify “northern-associated” and “southern-associated” alleles and ask whether there was an appreciable change between the two time points. To help reduce the role of linkage between sites, we further pared down SNPs so that no two SNPs were within 10kb of one another. This left 1671 examined SNPs. Notably, the northern-associated alleles increased in frequency by an average of 2.44%. Of the 1671 clinal SNPs examined, 62% of them (1036) had northern-associated alleles increase in frequency (binomial P < 0.00001 assuming a 50% null expectation). This pattern is in contrast to a random sample of non-clinal SNPs showing an average northern allele frequency shift of -0.11% (Figure 4). Based on this non-clinal SNP pattern, the observed genome-wide shift toward European ancestry between time points that we documented above does not seem to account for the imbalance of frequency changes at clinal SNPs. Instead, we hypothesize that outlier SNP ancestry clines have steepened in the last 35 years due to continued local adaptation at many of these clinal outlier loci.

**Figure 4.**
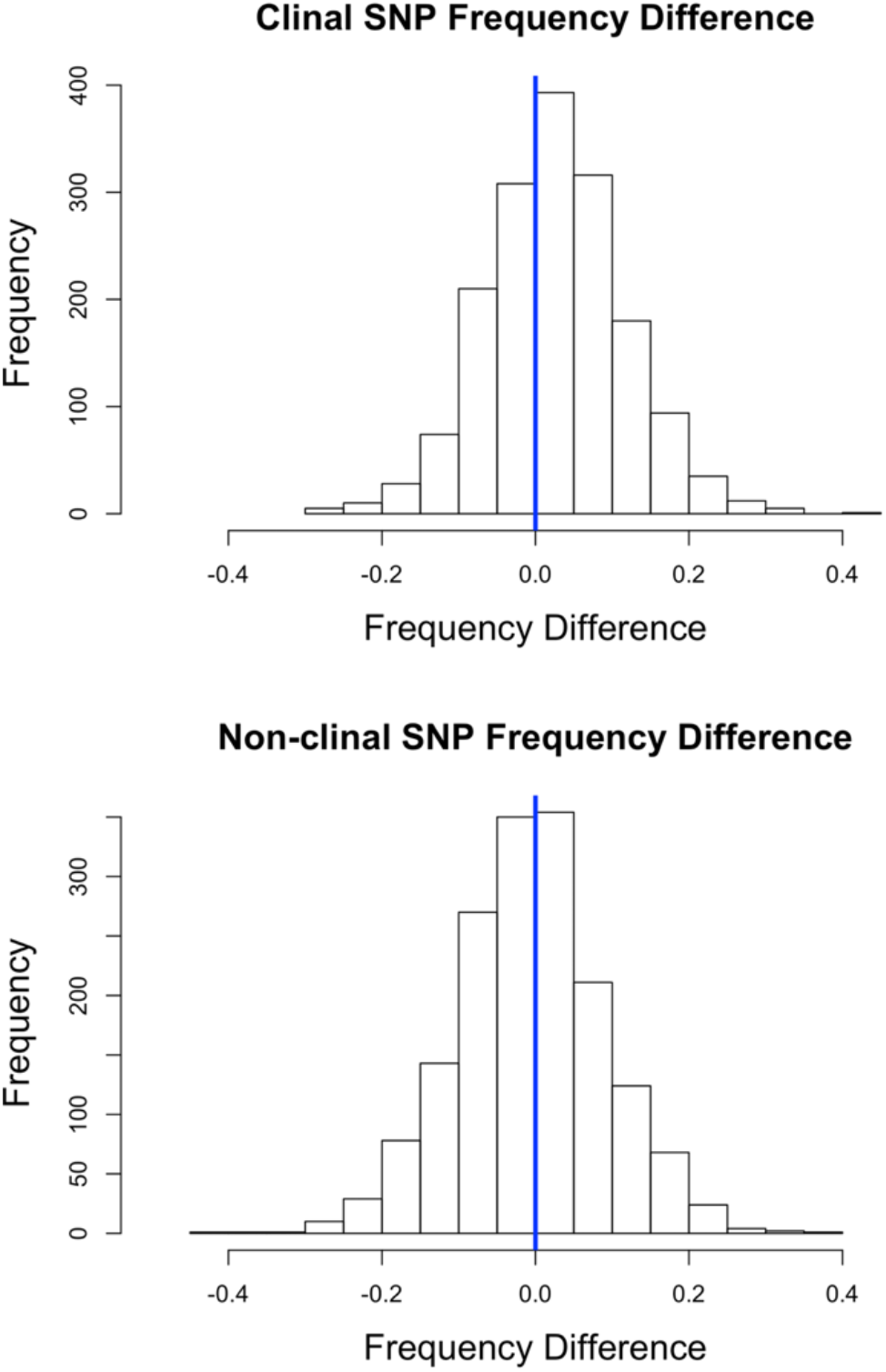
Northern-associated alleles at clinal outlier SNPs have tended to increase in frequency over time. The top histogram depicts 1,671 frequency differences of northern-associated alleles at clinal outlier SNPs. The bottom histogram depicts the frequency difference at 1671 SNPs chosen at random across the genome.

### Estimated Local Population Size is Relatively Small

Under neutralist assumptions, the long-term effective population size of *D. melanogaster* has been estimated to be on the order of 1,500,000 to 2,500,000 (Duchen 2013; Sprengelmeyer *et al*. 2020). However, one study suggested a much larger recent effective population size on the order of 10^8^ (Karasov *et al*. 2010). These population-scale sizes are likely much larger than local deme sizes, especially in a temperate environment with seasonal population size fluctuations. Smaller local census sizes have been estimated on the order of 1,000-10,000 (McInnis *et al*. 1982). It is important to have an accurate (or at least conservative) estimate of local population size to identify regions of recent directional selection that deviate from the predictions of genetic drift.

To estimate local population size, we simulated allele frequency trajectories of SNPs based on a simple Wright-Fisher model and fit the distribution of observed genome-wide frequency changes to distributions of simulated frequency changes corresponding to a given population size, while accounting for both individual and read sampling. Here we assumed that the bulk of the empirical distribution reflected neutral evolution over this short time period. We found that a population size of 9,500 individuals best recapitulated the empirical distribution of SNP frequency differences between the old and new population samples. Here, the empirical average magnitude shift in allele frequency was 0.0781 and the simulated standard deviation, with a population size of 9,500, was 0.0782.

Although the amount of genetic variation observed in *D. melanogaster* reveals that the long-term effective population size of the species is very large, our results suggest that a single local population may have distinct demographic dynamics. However, if natural selection is sufficiently widespread in the genome on short time-scales, it could bias our local *N*_*e*_ estimate downward. Such a bias would lead us to overestimate the strength of genetic drift, which is conservative with regard to identifying potential targets of recent natural selection.

### Potential Adaptive Differences Between Sampling Points

It has been shown that dozens of loci in the *D. melanogaster* genome exhibit evidence of seasonally-varying selection, with one allele becoming more common by spring and the alternate allele rising in frequency by fall (*e*.*g*. Bergland *et al*. 2014; Machado *et al*. 2021). We therefore wanted to separate seasonal allele frequency change from directional evolution across these ∼35 years. The latter signal can be isolated by the Population Branch Statistic (PBS; Yi *et al*. 2010), which uses three population samples to quantify genetic differentiation on one population’s lineage. Here, we applied PBS to a single old population sample (our focal population), the recent spring sample, and the recent fall population sample. PBS will therefore focus on genetic differentiation that separates the old population sample from both of the new samples, and the presence of two contrasting seasonal samples should serve to separate seasonal allele frequency evolution from the focal PBS lineage. In other words, at seasonally-oscillating SNPs where our new fall and spring samples show frequency differences, the old sample will have no branch-specific differentiation to inflate PBS unless its own frequency falls outside the range of the new fall and spring samples. We also accounted for inversion frequency differences between old and new population samples as described in the Materials and Methods.

We applied PBS to ∼5 kilobase windows (full results are provided in Table S7) and individual polymorphic sites along the genome (Figure 5A). We ran a Gene Ontology (GO) enrichment analysis on PBS outliers (the top 1% of windows) to identify functional categories that may hold adaptive differences between the sampling times (Figure 5B; Table S8). One of the top categories in this analysis was response to insecticide, which we discuss in more detail below.

**Figure 5:**
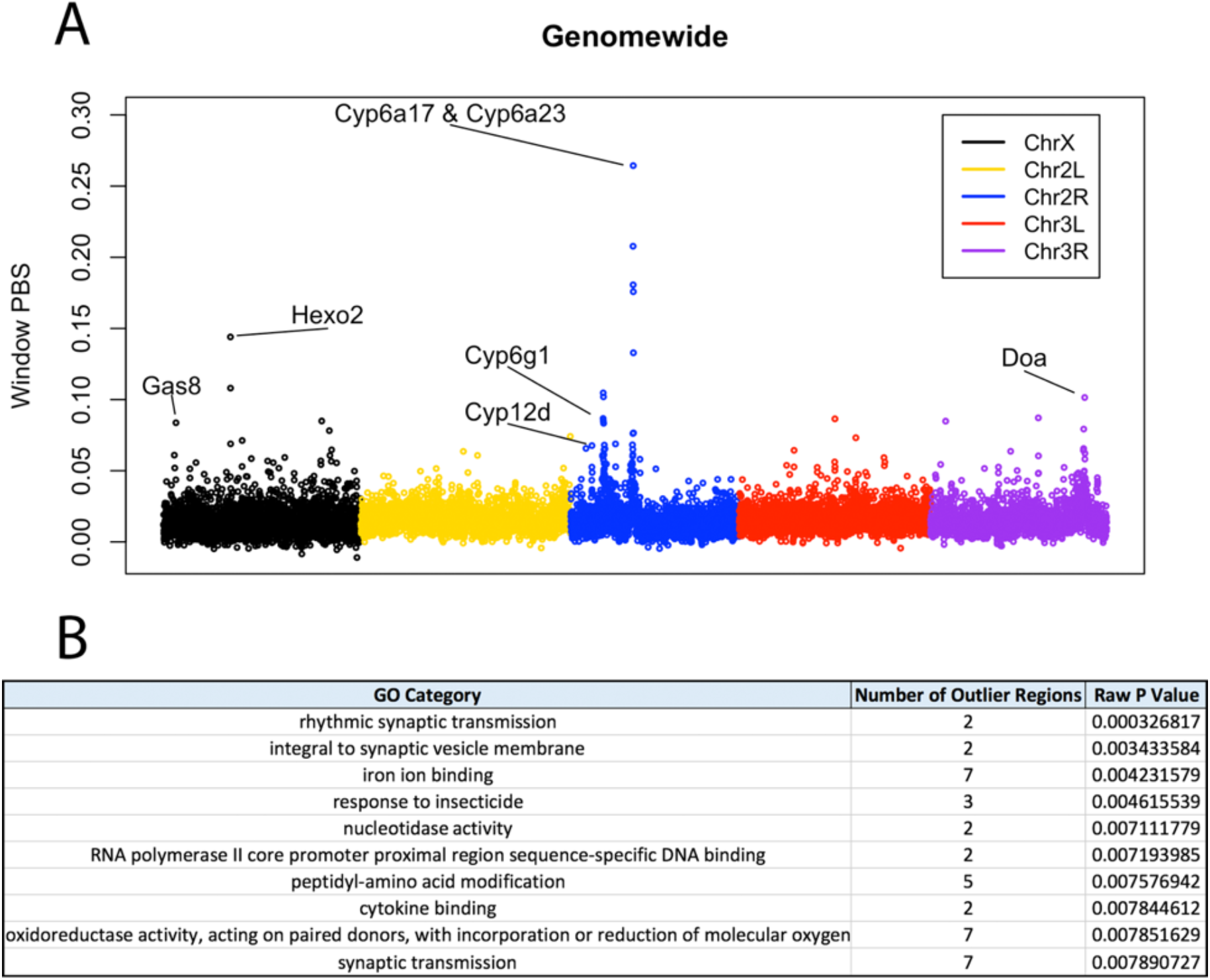
Population Branch Statistic and Gene Ontology Enrichment. (A) PBS at each window genome-wide. Gene names are discussed in the text. (B) The top 10 categories in our window PBS GO enrichment analysis.

Three other top categories were related to the nervous system. Nervous system GO categories have been enriched in other genomic scans for positive selection in *D. melanogaster* (*e*.*g*. Langley *et al*. 2012), including a study of parallel adaptation to cold climates (Pool *et al*. 2017).

### Insecticide Resistance as a Likely Target of Selection

We used simulation to assign a P value to every empirical window PBS and also to the highest SNP value of PBS within the window (max-SNP PBS). Empirical windows were divided into five bins based on recombination rates (Comeron *et al*. 2012), and 2.5 million simulations were run for each bin. Briefly, we simulated the species demography from Sprengelmeyer *et al*. (2020). This demography consists of 9 populations sampled throughout Africa and Europe. To include the North American Rhode Island population, we added a 10th population consisting of an admixed population of Cameroon and France migrants. The simulated admixture proportions reflected our estimated genome-wide African and European ancestry levels from our old sample. Each window was then assigned a raw P value based on the proportion of neutral simulated replicates from its recombination rate bin that yielded a greater window or max-SNP PBS value.

In Table 1, we display our top outlier regions. Our strongest genome-wide outlier region had a window-PBS value of 0.278 and spanned 78 windows. The P value associated with the top window in this region was below a Bonferroni-corrected genome-wide significance threshold (*i*.*e*. 0.05 divided by the total number of windows). Notably, this window contained a pair of cytochrome P450 genes, *Cyp6a17* and *Cyp6a23*. Previous research has identified derived alleles in which only one chimeric gene is present, and these chimeric (primarily *Cyp6a23*-derived) alleles were found to segregate at high frequency in the DGRP population (Good *et al*. 2014). Later research showed that disruption of *Cyp6a17* confers resistance to the deltamethrin class of insecticides in *D. melanogaster* (Battlay *et al*. 2018), and that this locus shows the strongest association with deltamethrin resistance among DGRP strains genome-wide (Battlay *et al*. 2018; Duneau *et al*. 2018).

**Table 1.**
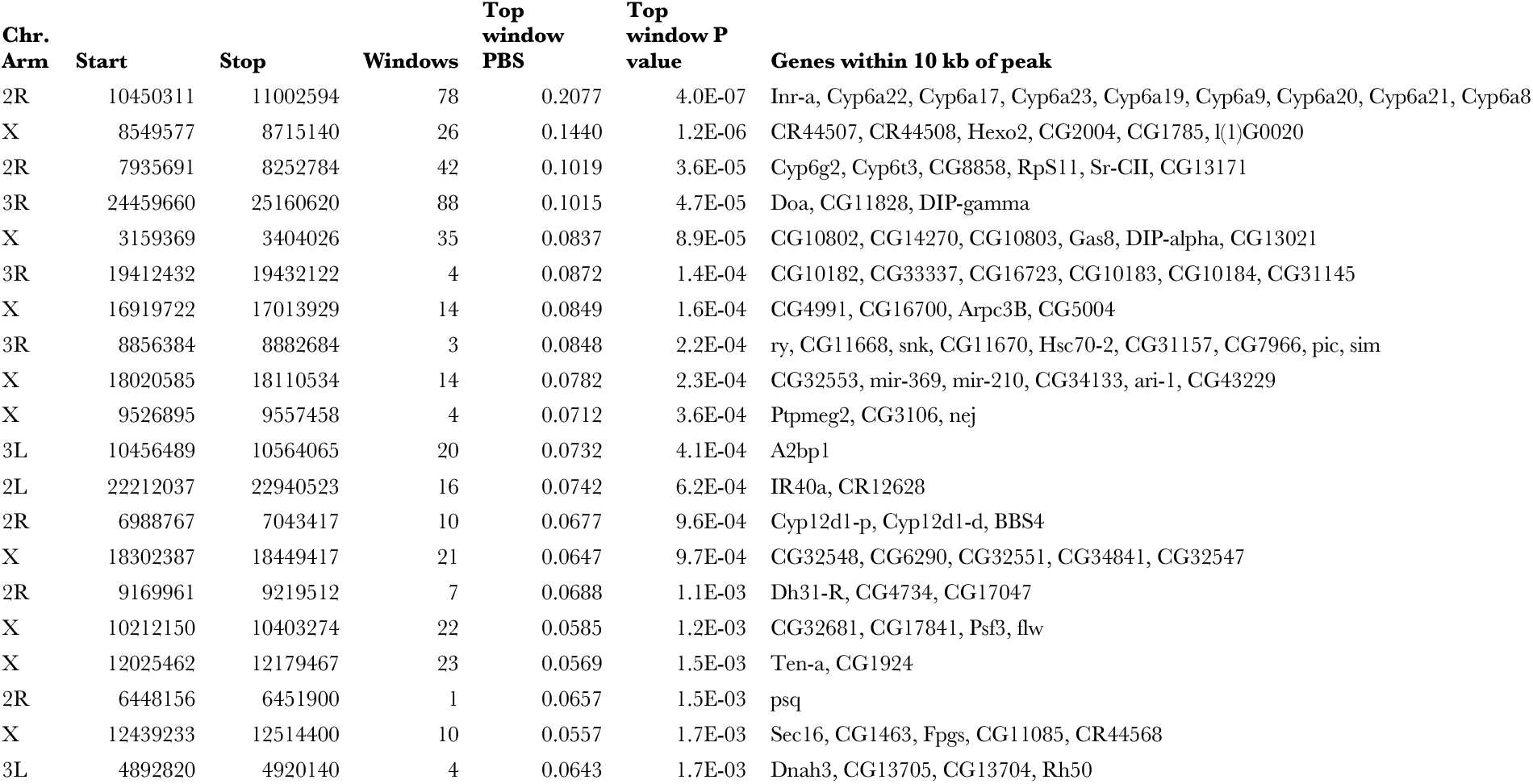
Outlier regions containing the top Population Branch Statistic values genome-wide, indicating candidates for temporal evolution. Regions were defined as described in the Materials and Methods. Raw P values based on demographic simulations are also provided. Coordinates reflect release 5 of the *D. melanogaster* reference genome.

Given the clear signal of genetic differentiation we observed at this locus, we estimated the frequency of the intact *Cyp6a17* + *Cyp6a23* allele in both our old and new samples. On average, we observed this “intact allele” at 23.53% frequency in the old data versus 51.08% in the new data. We also examined the frequency of the intact allele in each of the sampling years of the old data, which revealed a striking result. We found that the frequency of the intact allele increased over the 8 year period in which the older population sample was collected, rising from 0% in 1975 to >50% in 1983 (Figure 6A), possibly revealing a selective sweep in action. This case curiously mirrors that of the olfactory receptors *Or22a* and *Or22b*, in which selection also appears to have recently acted in favor of an ancestral two-paralog haplotype in some populations, at the expense of a derived deletion variant carrying a single chimeric gene (Aguadé 2009; Mansourian *et al*. 2018).

**Figure 6:**
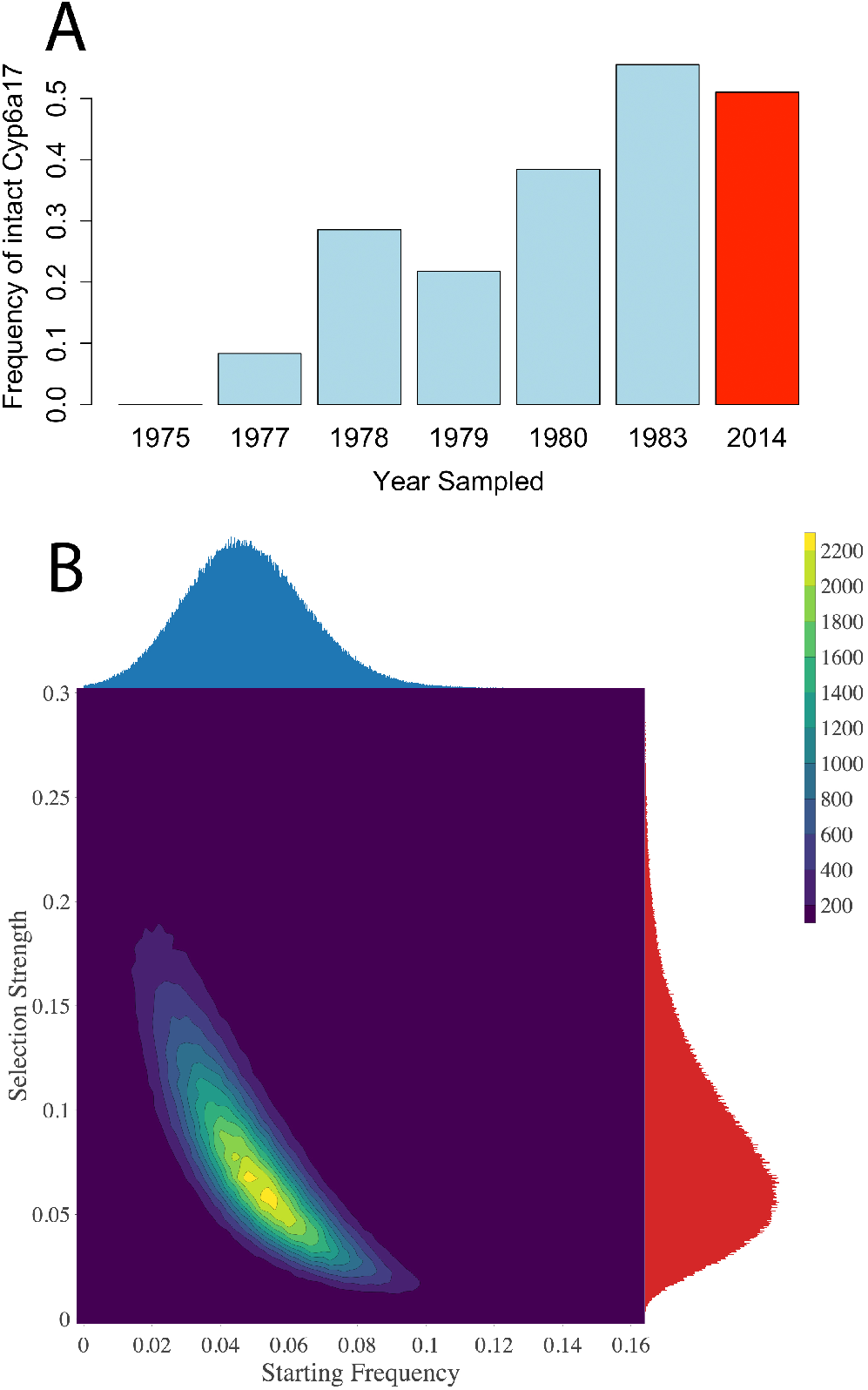
The frequency of the intact *Cyp6a17* + *Cyp6a23* allele shifted rapidly under strong selection. (A) The frequencies of the intact allele in each of the 6 sampling years and an estimate of the frequency in modern populations are shown. (B) Results of an ABC analysis to infer the selection strength and starting frequency of the intact allele that best recapitulate our empirical sampling results between 1975 and 1983.

By examining coverage at *Cyp6a17* in our seasonal samples, we estimated that the intact allele is still only around 50% frequency in modern Providence populations. Thus, after its rapid rise, it appears that the frequency of an intact *Cyp6a17* did not meaningfully change between 1983 and 2014. These patterns might reflect changes in insecticide usage with time, or might potentially represent the impact of balancing selection due to fitness costs associated with insecticide resistance, a well-studied phenomenon in insects (Kliot and Ghanim 2012). It is not clear that any cost of resistance should be dramatic enough to impact fixation probabilities in small laboratory cultures (and hence frequency estimates in our old population sample), but we can not formally exclude this possibility.

We performed approximate Bayesian computation to investigate the strength of natural selection required to observe the empirical frequency shift over eight years. We simulated a simple Wright-Fisher sampling over an eight-year period, assuming 15 generations per year (Turelli and Hoffmann 1995; Pool 2015), and sampled to match our empirical counts. We varied the starting allele frequency and the selection strength, and rejected any simulation that did not exactly match empirical counts of the sampled allele. We obtained maximum a posteriori (MAP) estimates (univariate modes of 0.1% bin sizes) of 6.75% for selection strength (95% CI 2.10% to 19.02%) and 4.55% for initial frequency (95% CI 1.49% to 8.47%), as further depicted in Figure 6B. Although the details of natural selection at this locus may have departed from the constant additive advantage simulated here (as discussed above), it does seem clear that strong selection and an appreciable starting frequency are required to explain the observed results. A relatively high initial frequency of this beneficial allele is perhaps not surprising since it appears to reflect the ancestral arrangement (Good *et al*. 2014) as opposed to a derived variant entering the population at very low frequency.

Another known insecticide resistance gene, *Cyp6g1*, was contained within our third-highest outlier region. This gene has been shown to confer resistance to dichlorodiphenyltrichloroethane (DDT) and other insecticides via the insertion of a transposon upstream of the transcription start site (Daborn *et al*. 2002; Chung 2007; Battlay *et al*. 2016). It has also been shown that ongoing selection at this locus is caused by a duplication and additional transposable element insertions at this locus (Schmidt *et al*. 2010). Our outlier region spanned 42 windows, and the top window had a PBS value of 0.102. The P value assigned to this window (p=3.64E-5) was just above the Bonferroni-corrected significance threshold. This window was not directly over *Cyp6g1*, instead lying about 27 kb downstream, and 10 kb upstream of the closely related gene *Cyp6g2*, which has also been shown to confer insecticide resistance (Daborn *et al*. 2007).

A third well-known locus that confers resistance to insecticides was also amongst our top 20 regions genome-wide. This region spanned 10 windows, and the top window included *Cyp12d1-p* and *Cyp12d1-d*. The PBS value at this window was 0.0677 and its P value was 0.00096. When mapped reads were viewed with IGV (Thorvaldsdóttir et al. 2013), few to no reads mapped uniquely to the *Cyp12d1-p* and *Cyp12d1-d* locus at either time point, potentially due to high sequence identity between them. Instead, we observed high PBS SNPs flanking the locus. Gene copy number variation at this locus is known to remain polymorphic in natural populations, and it has been linked to resistance to xenobiotics including caffeine (Najarro *et al*. 2015), although it was not found to correlate with DDT resistance (Schmidt *et al*. 2010). However, *Cyp12d1* is inducible by DDT and its overexpression confers DDT resistance (Daborn *et al*. 2007). *Cyp12d1* expression level was also found to correlate with malathion resistance among DGRP strains (Battlay *et al*. 2018). Notably, a deletion spanning both *Cyp12d1* genes was found to show parallel frequency clines with latitude in North American and Australia, with the intact alleles more common in higher latitude populations (Schrider *et al*. 2016). Hence, our high PBS SNPs may be tracking linked copy number changes at this locus.

Another well-known insecticide resistance gene, *Acetylcholine esterase* (*Ace*), offers resistance to organophosphates and carbamate insecticides (Mutero *et al*. 1994). Windows associated with this gene had a PBS value in the top 1% genome-wide. Known insecticide resistance mutations in this gene have been previously reported (Mutero *et al*. 1994, Menozzi *et al*. 2004, Karasov *et al*. 2010). A set of three mutations within twenty base pairs of each other make up a resistant haplotype. All three of these resistance alleles segregated at low frequency in the old samples (F330Y at 0%, G265A at 2.08%, and I161 at 16%) and segregated at 29%, 29%, and 41% respectively in the new, seasonal samples. Thus, over this relatively short time period, these three mutations associated with insecticide resistance increased by an average of 26.97%.

Our data suggest that insecticides have been a major driver of evolution in the *D. melanogaster* genome in North America over the last 35 years. Interestingly, all four insecticide loci described here have recently been identified as candidate regions of recent adaptive introgression into African *D. melanogaster* populations from non-African sources (Svedberg *et al*. 2020), underscoring the likely importance of insecticide evolution in admixed populations. Additionally, the temporal nature of our data can reveal important information about the trajectory of an allele and its underlying selection coefficient, allowing researchers to more accurately model the genetic basis of adaptation in human-associated insect species including crop pests and invasive species.

### Male Reproduction as a Potential Target of Natural Selection

It is possible that male reproductive performance may have also been an important evolutionary target between our time points. At least three genes in our top 20 PBS outliers may play important roles in male mating success. Our second highest PBS outlier region was on chromosome X and covered 26 windows. The top window in this region had a window PBS of 0.144 and a P value of 1.2E-6, which was below the Bonferroni-corrected significance threshold. This window contained two pseudogenes as well as *Hexosaminidase 2* (*Hexo2*). Zooming in to the SNP level of this region, we observed a collection of high PBS SNPs as well as a modest PBS peak in the intergenic regions between *Hexo2* and the pseudogenes (Figure S2A). The gene product of *Hexo2* is found in the plasma membrane of sperm in *D. melanogaster* (Cattaneo *et al*. 2006) and has a possible role in fertilization and sperm-egg interactions (Intra *et al*. 2017).

Two other genes in our top 20 regions have possible effects on reproduction as well. The first, *Darkener of apricot* (*Doa*), was within the top window of our fifth highest outlier region. The PBS at this window was 0.101 and had a P value of 4.68E-5. *Doa* spanned four windows, though the PBS signal appeared to localize toward the end of the gene (Figure S2B). Researchers have used artificial selection experiments to show that this gene affects aggression in flies (Edwards *et al*. 2006, Zwarts *et al*. 2011). A subsequent study demonstrated that mutations at *Doa* disrupt sex-specific splicing of *doublesex* pre-mRNA, resulting in the feminization of male cuticular hydrocarbon profiles and the masculinization of female cuticular hydrocarbon profiles, along with disruption of associated courtship behavior (Fumey and Thomas 2017). A third gene that affects fertility was also within our top outlier regions. *Growth arrest specific protein 8* (*gas8*) spanned the top two windows in our fifth highest outlier region. This gene has been associated with sperm fertility in mice (Yeh *et al*. 2002). In *D. melanogaster*, knockdown of this gene causes infertility in males (Zur Lage *et al*. 2019). However, male reproduction functions were not enriched in our GO analysis cited above, and further studies are needed to determine whether the recent evolution of genes with roles in male reproduction is associated with ongoing male-male reproductive competition, male-female reproductive coevolution, or the optimization of male reproductive success in local or changing environments.

### Other Targets of Interest

A region on chromosome X was our 7^th^ highest PBS outlier (PBS=0.0849, P value 0.000156). Zooming in on the window revealed a SNP pattern that localized over two genes, *CG4991* and *CG16700* (Figure S2C). This region has previously been identified as a target of positive selection between African and non-African populations (Svetec *et al*. 2011), likely due to cold tolerance adaptations (Ayroles *et al*. 2009, Wilches *et al*. 2014).

### A Complementary Scan to Confirm Window Outliers and Identify SNP-level Outliers

Because we were comparing two distinct types of sequencing data (individual sequences and pool sequences), we wanted to complement our original scan with a scan where all datasets went through the same pipeline. We down-sampled raw reads from our 64 old isofemale line genomes so that they all had the same number of reads. We then combined the reads to emulate a pooled set of sequences and performed a PBS scan with this “pseudo-pool” set. The PBS outliers at the window level were highly consistent across the two scans. All outliers in Table 1 were also outliers in the “pseudo-pool” scan. Hence, the results described above seem robust to differences in sequencing strategy between samples.

Especially in the case of soft sweeps, it is possible that some positive selection signals spanned less than a full window and were missed by the window scan described above. However, when we examined top outliers for maximum SNP PBS (the highest SNP PBS value within a given window) from our primary scan that were not also window PBS outliers, we found that these SNP-specific outliers were not well-replicated in our pseudo-pool analysis (results not shown). Because these outliers could reflect artifacts driven by differences in data processing between pool and individual genomes, we did not examine them individually. Instead, we focused our SNP-oriented analysis solely on the pseudo-pool scan outliers for max-SNP PBS, since these are not impacted by any differences in data processing.

In Table 2, we present 14 windows where the maximum SNP PBS value was in the top 1% genome-wide in our initial scan, and also in the top 2.5% genome-wide in the complementary pseudo-pool scan, while the window PBS value was a non-outlier (in the *bottom* 95% genome-wide) in the original scan. The top SNP PBS signal in this group was from the gene *heavyweight* (*hwt*) (Figure S3A). Variation at this gene is associated with body mass among DGRP strains (Nelson *et al*. 2016). Both *Neurospecific receptor kinase* (*Nrk*; associated with the second-highest SNP PBS signal; Figure S3B) and *Dystrophin* (*Dys*; Figure S3C) also contained a high SNP PBS but lacked a window signal. These two genes interact genetically to control neuron behavior in the eye (Marrone *et al*. 2011), and echo the potential nervous system evolution suggested by our GO enrichment analysis of window PBS outliers. Panels D-F in Figure S3 display SNP patterns at three other regions where there is a relatively strong SNP signal but weak window signal.

**Table 2.**
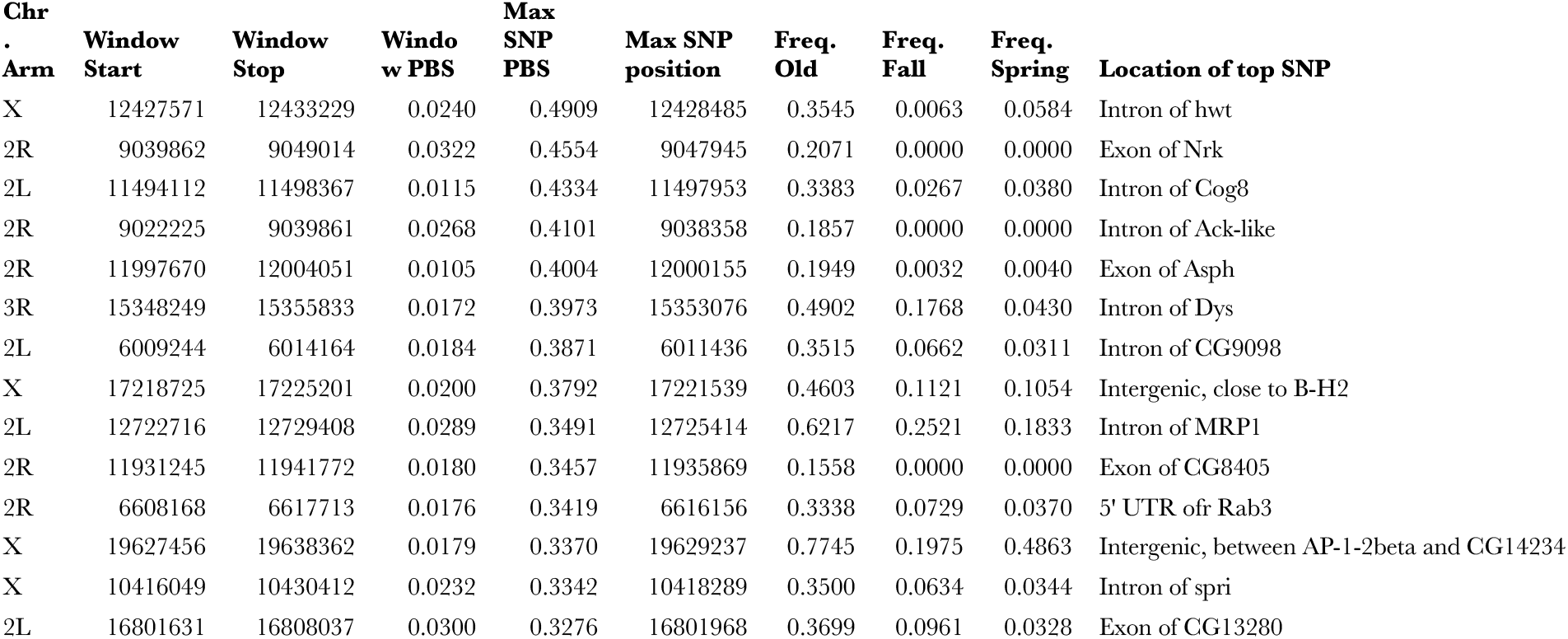
Windows containing the highest SNP PBS values genome-wide are shown, excluding those associated with window PBS outlier regions, based on our “pseudo-pool” treatment of the old sample data. Coordinates reflect release 5 of the *D. melanogaster* reference genome.

### Genome-wide Enrichment of PBS Outliers

Above, we described the generation of a *P* value for every window and max-SNP PBS value based on demographic simulations. Under this null hypothesis, we would expect our neutral simulations to recapitulate our empirical distributions of window PBS and max-SNP PBS, yielding a uniform distribution of P values. Instead, we saw a slight enrichment of low P values for the window PBS statistic, and a similar enrichment for the max-SNP PBS statistic (even when focusing only on outliers confirmed by the pseudo-pool scan; Figure 7). Based on these observed outlier enrichments, we asked how many inclusively-delimited outlier regions would need to be removed before the enrichment disappeared (see Materials and Methods).

**Figure 7:**
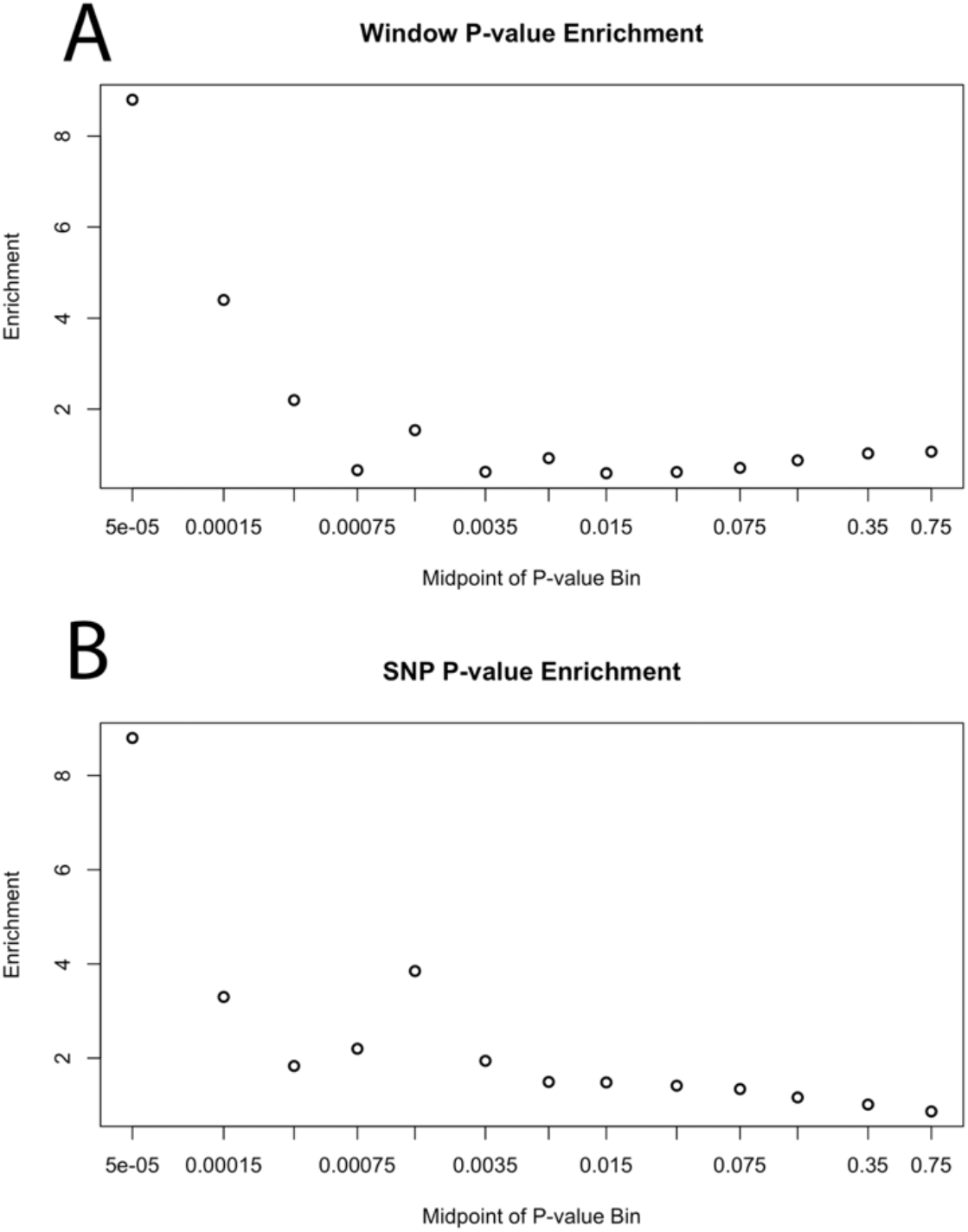
A genome-wide enrichment of elevated genetic differentiation between old and new population samples was observed. The enrichment of low raw P values, indicating higher PBS values than expected under the neutral demographic model, is depicted for (A) window PBS, and (B) maximum SNP PBS per window. As noted in the main text, the window enrichment can be accounted for by just a few broad outlier regions, whereas ∼20 SNP PBS outlier regions must be removed to account for the latter enrichment.

For window PBS, we found that only the top 4 regions needed to be removed to account for the observed enrichment of outlier PBS values compared to the neutral model. In contrast, for max-SNP PBS 20 regions were removed with this deterministic approach, accounting for 4.83% of the genome. We complemented this max-SNP PBS analysis with a random outlier region removal approach, in which an average of 17 regions were removed, accounting for 3.87% of the genome. These results suggest that roughly 20 loci experienced strong enough selection between our time points to contribute to the observed genome-wide excess of PBS outliers, and that relatively more targets of strong selection may affect genetic differentiation at the SNP level than the window level.

## Discussion

In this study, we have compared genomic variation of recently collected samples of *Drosophila melanogaster* to collections made approximately 35 years prior. This unique dataset has given us insight into a range of population genetic phenomena, including local effective population sizes, inversion frequency dynamics, and the potential targets and apparent prevalence of relatively strong natural selection.

Our results point to aspects of fine-scale demography that remain poorly understood, even in this population genetics model system. We obtained an estimate of local effective population size (9,500) that is more than two orders of magnitude lower than long term population genetic estimates (*e*.*g*. Sprengelmeyer *et al*. 2020), and at least four orders of magnitude lower than suggested from a study of recurrent adaptive mutations (Karasov *et al*. 2010). While these estimates are not necessarily in conflict, and the potential contribution of natural selection to differences among them is unclear, their contrasting magnitudes nevertheless highlight the need for further study of local population dynamics in this species. Seasonal bottlenecks likely play some role in this finding, especially given that effective population size across time is equal to the harmonic mean of generation-specific population sizes. Further, our findings regarding inversion frequencies differing among spring but not fall sampling sites may point to the interplay between seasonal bottlenecks and gene flow that may later homogenize such local effects. More detailed models of local deme size, seasonal population size fluctuations, and connectivity among demes would enhance the null models that temporal population genomic data sets such as ours can be compared against. Hence, we suggest that a clearer understanding of local population dynamics across diverse environments in this species should be pursued.

When the results of a population genomic scan for natural selection are compared against a demographic model at all, it is rare for any outlier to show a stronger departure from the null model than would be expected at any locus in the genome by chance. In our study, 3 outlier regions met this rather conservative threshold. Among these extreme outliers was the *Cyp6a17*/*Cyp6a23* locus, for which we were able to estimate parameters of positive selection with greater precision than typically obtained from a single snapshot of genetic variation. These results highlight the potential of temporal samples from natural populations to strengthen population genetic inferences.

The notably stronger enrichment of max-SNP PBS outliers versus window PBS mirrors the results of Pool et al. (2017) that SNP-level differentiation was more sensitive in detecting evidence of parallel adaptation to cold environments. Soft sweeps, in which a selected variant was already present on multiple haplotypes, might be expected to produce stronger SNP than window patterns of genetic differentiation. Selection on standing genetic variation is highly plausible in light of this species’ high genetic diversity and relatively recent worldwide expansion (*e*.*g*. Sprengelmeyer *et al*. 2020), and recurrent mutation may be relevant to its ongoing adaptation as well (Karasov *et al*. 2010). Further work is needed to formally identify the evolutionary scenarios in which SNP-level analyses of genetic differentiation are critical.

Our results also suggest that the integration of temporal and spatial population genomic data may be especially fruitful. A key example is our finding that for SNPs previously shown to show unusually strong latitude clines in allele frequency, the “northern” allele has tended to increase over the past ∼35 years in the relatively cool environment of this northeast US population. Notably, this increased frequency of “northern” alleles has occurred in spite of a roughly 1°C local temperature increase during this period (Lenssen et al. 2019; GISTEMP Team 2021). These results suggest that local adaptation along the Atlantic coast of the US may be an ongoing process. Our finding of a genome-wide increase in the frequencies of alleles with European as opposed to African origin is compatible with that hypothesis as well, but further data and analysis are needed to more fully understand the contributions of directional selection, epistatic selection, and population history in admixed North American populations of *D. melanogaster*. Expanded spatiotemporal population genomic sampling, including that already being conducted on shorter time scales (Kapun *et al*. 2020, 2021; Machado *et al*. 2021), may further leverage the complementary spatial and temporal signals of population genetic processes.

The increasing availability of temporal population genomic data may motivate further studies developing novel methodology to estimate the relative roles of neutral and non-neutral processes in shaping genomic variation. As an example, Buffalo and Coop (2020) developed a method to investigate the genome-wide influence of natural selection based the on temporal covariance of allele frequency changes (i.e. a consistent direction of change across multiple sampling time points), and applied it to experimental evolution data. Given the data we have already collected from this Rhode Island *Drosophila* population, collecting additional genomic data in future years may allow such a method to be applied to this natural population.

Extending the temporal scale of population genomics in natural populations should hold increasing importance in evolutionary biology. Such studies may not always be limited to contemporary sampling efforts: previously collected and sequenced population samples may be potential targets for resampling from current populations, and even museum collections may be potential sources of population genomic data (Mikheyev *et al*. 2015). Our study provides one example of an analysis framework that allows allele frequency change within a specific time interval to be isolated. Especially in rapidly reproducing organisms like insects, temporal population genomic studies may play increasingly important roles in monitoring adaptation to climate change, pesticide use, and other environmental alterations, with important implications for agriculture and conservation in addition to basic science.

## Materials and Methods

### Fly Collection

The 65 isofemale lines collected between 1975 and 1983 were originally collected by Dr. Margaret Kidwell and had been maintained at the University of Wisconsin by Dr. Rayla Temin. These strains had been maintained in vials, in a separate tray from other stocks. Although it is possible for contamination to occur in fly stocks over time (*e*.*g*. Frochaux *et al*. 2020), and we can not be certain that none has occurred in our strains, it seems clear that contamination would need to affect a large fraction of our strains in order to meaningfully impact analyses such as the frequency-based outlier scan described below. No phenotypes consistent with common laboratory mutant stocks (*e*.*g*. eye color, body color, wing morphology) were observed in any of them. Additional evidence against contamination of these stocks was obtained from the IBD and PCA analyses described below.

We note that the small population sizes of laboratory fly cultures are not conducive to natural selection unless it is extremely strong. There is also a limited and diminishing amount of genetic variation that goes into an isofemale line, and the population mutation rate in a lab culture is also low, such that parallel mutations across multiple lines are very unlikely. If we assume that the rate of mutation to a specific nucleotide is 1.7e-9 (Huang et al. 2016), and that there were 910 generations between collection and sequencing (conservatively assuming a generation every two weeks in the lab across 35 years), then the binomial probabililty that a given new mutation from one of our 64 lines is also sampled from even one additional line is only 1%. New mutations should therefore have little effect on allele frequencies across the full population sample. Hence, we consider these living strains to be a good proxy for genetic variation in the source population at the time of sampling, with particularly little opportunity for sample-wide shifts in allele frequency between collection and present.

For the newer collections, flies were collected from 5 trap locations in Providence, Rhode Island in Fall 2014 and Spring 2015. These locations (Weymouth Street, Brown Street Park Community Garden, Center for Environmental Studies Garden, Student Garden, and Fox Point Community Garden at Gano Park) centered around the original sampling location (Weymouth Street) and none were more than 1.7 km apart. In the fall samples, the first four pools contained 41 flies each from the Brown Street, Environmental Studies, Student Gardent, and Fox Point sites. The fifth pool contained 41 flies from mixed locations. A sixth pool contained 42 F1 female offspring from 6 wild-caught females from the Weymouth Street site. When analyzing allele frequencies, each pool was weighted based on effective pool size as described below. The spring samples consisted of 12 total pools, each with 34 flies.

### Genomic Sequence Data Collection

A separate library was prepared for each of the 65 old individual strains (from 30 adult females of each strain) and from the 18 new fall and spring pools. Mean sequencing depth per individual genome ranged from 5.26X to 71.79X. 100 base pair reads were aligned as described in Lack *et al*. (2015), except with a single round of mapping to the *D. melanogaster* (v5.57) reference genome instead of a second round of mapping to a genome-specific reference genome. We chose a single round of mapping to more closely align with the single round of mapping that the pooled sequences necessarily will go through. Briefly, reads were aligned using BWA aln v0.5.9 (Li and Durbin 2010), and unaligned reads were then mapped with Stampy v1.0.20 (Lunter and Goodson 2011). All reads with a mapping quality score below 20 were discarded. We then used Picard version v1.79 (http://picard.sourceforge.net/) to sort the alignment by coordinates and remove optical duplicates. Assemblies were improved around InDels using the GATK v3.2 InDel Realigner (Depristo *et al*. 2010, McKenna *et al*. 2010;). Consensus sequences for homozygous heterozygous regions (see below) were generated using GATK haploid and diploid consensus sequence calling, respectively, as described in Lack *et al*. (2015). This pipeline was used to maximize the comparability of our new individual strain genomes to data from the *Drosophila* Genome Nexus (Lack *et al*. 2015, 2016).

To generate the “pseudo-pool” dataset, we down-sampled each of the 64 bam files from the individual sequences dataset so that each strain had an equal number of aligned reads. We did the downsampling with the command line “samtools view -s fraction -b data.bam > downsampled_data.bam” where “fraction” was the proportion of aligned reads kept. We then merged all 64 downsampled bam files using samtools and generated a pileup and subsequent sync file as we did with our other pool sequences.

Mean sequencing depth for the pools are provided in Table S2. Reads were aligned using the same pipeline as the individual genomes up through InDel realignment. Following InDel realignment, pileup files were then generated using Samtools v1.3.1, and sync files were generated using PoPoolation2 v1.201 (Kofler *et al*. 2011), requiring a minimum quality score of 20.

### Heterozygosity

Residual heterozygosity often persists in fly stocks even after many generations of full-sibling mating. It is important to identify such regions of heterozygosity, since a putatively heterozygous region constitutes two random allele draws from a population. We thus sought to identify such regions of heterozygosity in order to identify correct sample sizes at any given site in the genome. We applied a hidden Markov model (https://github.com/russcd/Heterozygosity_HMM) to annotate inbred and outbred regions in the partially inbred old samples. On average, 32.6% of the the old isofemale line genomes were called as heterozygous (compared to 13% for the strongly inbred DGRP; Lack et al. 2016). Full results are provided in Table S9.

### Quality Assurance Checks (IBD, PCA)

Relatedness between sampled individuals violates assumptions of many population genomic models. Such relatedness could result from initial sampling of related individuals or from mishandling or mislabeling of lab samples in the last 35 years. To identify such instances of identity by descent (IBD), all pairwise comparisons between old samples were made. We also included the reference genome sequence in this analysis to guard against contamination from a common laboratory background. Chromosomes were compared in 500 kb windows sliding in 100 kb increments. Any window with fewer than 0.0005 pairwise differences per site was considered putatively IBD. Some genomic intervals (e.g., centromeric regions) exhibit large scale IBD across populations, suggesting explanations other than relatedness. These regions of recurrent sequence identity were not permitted to seed new IBD regions, but neighboring IBD regions were allowed to extend through them. We identified instances of “relatedness IBD” between two genomes when genome-wide IBD tracts exceeded 5 Mb. Such instances were masked to ‘N’ for one of the two genomes. This IBD threshold will limit our detection to pairwise comparisons of homozygous regions, and hence some IBD may persist elsewhere. However, given the relative rarity of IBD between pairwise homozygous regions, any residual IBD should have limited impact on our analyses.

We applied principal components analysis (PCA) to each major chromosome arm of the old samples to identify any strain with aberrant divergence. We also included several African populations (10 Cameroon strains, 10 Gabon, 6 Nigeria, 5 Guinea), 98 France strains, and two North American populations (19 from Ithaca, New York, 131 from Raleigh, North Carolina). Putatively heterozygous regions in all strains were masked for this PCA analysis. See methods of Lack *et al*. (2015) for masking heterozygosity in the non-Rhode Island genomes, and see the “Heterozygosity” subsection above regarding identifying heterozygosity in the old Rhode Island strain genomes. We used SNPRelate release 3.9 (Zheng *et al*. 2012) for the PCA analysis.

### Effective Numbers of Sampled Individuals and Alleles from Pool-seq Data

One of the major drawbacks of utilizing pooled sequencing is that we cannot assume that every read is a random draw from a population, due to the dual binomial sampling processes of individual and read sampling. Though allele frequency estimates are accurate in a large enough sample (Gautier *et al*. 2013), nominal sample sizes may be misleading if individuals contribute

DNA unequally to a genomic library. Because sample size is an important parameter when estimating *F*_*ST*_, we must take measures to obtain an unbiased estimate of the effective pool size, *n*_*e*_.

We applied the method introduced in Gautier *et al*. (2013) to estimate the effective pool size, defined as the number of equally contributing diploid individuals in a pool. Our fall data contains 5 pools of 41 individuals from 5 distinct sampling locations and a 6^th^ pool containing 42 flies (which had 7 F1 offspring from each of 6 wild-caught females). Here, we are treating pools from the same season as replicates, and while they might have some structure between them, any true allele frequency differentiation would downwardly bias our estimates of effective pool size, which is conservative for our analyses.

The effective pool size method estimated no error in the first 5 pools (*n*_*e*_=41for all pools) and an *n*_*e*_=10for the 6^th^ pool. The 6^th^ pool offered a good control of the method, as we expected this pool to have a lower *n*_*e*_ due to the relatedness of individuals in the pool. The method suggested a lower effective pool size number in the spring data. Here, each pool consisted of 34 flies. The effective pool size ranged from *n*_*e*_ = 23 to *n*_*e*_ = 34 with a median of *n*_*e*_ = 31.

We could then use the method of Ferretti *et al*. (2013) to estimate sample size at a given site. Here, we wish to explicitly estimate the number of *j* unique lineages sampled at a site given *n*_*r*_ reads and *n*_*c*_ equally contributing chromosomes in a pool. Ferretti *et al*. (2013) showed that 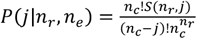, where *S*(*n*_*r*_ *j*) are the Stirling numbers of the second kind, defined as the number of ways to partition *n*_*r*_ reads into *j* nonempty sets. For our diploid data, we set *n*_*c*_ = 2 *n*_*e*_. We used the above formula to generate a lookup table giving the expected number of lineages for each potential number of sampled reads and effective pool size, by applying the expectation formula 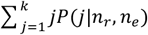, using the R package GMP v0.5-13.6. This procedure thus provided estimates of the expected chromosomal sample size for each site (*i*.*e*. the estimated number of natural alleles sampled).

### Ancestry

We implemented a hidden Markov model (Corbett-Detig and Nielsen 2017) to estimate the proportion of African versus European ancestry. This method is general and can be used on individual genomes or on high ploidy data (i.e., pooled data). It utilizes short read pileup data to model ancestry across the genome as a function of sample allele frequencies within an admixed population. The maximum likelihood probability of each ancestry state at each panel SNP is output. For a diploid genome, for example, maximum likelihood states for homozygous European, heterozygous European and African, and homozygous African are output. This method also allows for variable ploidy across the genome to account for partially inbred chromosomes, allowing us to model inbred segments as a single haploid chromosome and outbred segments as a diploid chromosome individually in each old genome. We generated ploidy maps as described above. For the high ploidy datasets (pooled data), maximum likelihood probabilities are output for each of ploidy + 1 states.

### Identification of Common Inversions

We implemented the method introduced in Kapun *et al*. (2014) to estimate inversion frequencies in each of the 18 pooled samples. For each studied inversion, this method required a set of fixed differences between inverted and non-inverted karyotypes. We used the inversion-specific markers given in Table S4 in Kapun *et al*. (2014). To estimate the inversion frequency in an individual pool, we calculated the average frequency across all inversion-associated SNPs whose coverage exceeded a minimum of 10 reads. Seasonal inversion estimates were made by weighting pools by the estimated effective pool size (*n*_*e*_), such that 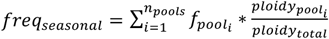. Here, *n*_*pools*_ is the total number of pools in a given season, 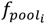 is the average frequency across inversion-associated SNPs in the *i*^*th*^ pool, 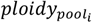 is the effective pool size in the *i*^*th*^ pool, and *polidy*_*total*_ is the sum of effective pool sizes across all pools.

To determine inversion status of the individual genomes in the old sample, we examined the diploid calls at each inversion-associated SNP described above. These diploid calls fell into two classes: most of the SNPs were homozygous for the non-inversion SNPs or most of the SNPs were heterozygous for the inversion SNPs. The former we classified as free of inversion and the latter we classified as heterozygous for the inversion. The inversion frequencies in the old genomes were calculated as the number of inversion heterozygotes divided by twice the total number of genomes in the dataset.

### Frequency Differences in Common Inversions

There is some evidence to suggest that inversion frequencies differ seasonally in *Drosophila melanogaster*. Our dataset provided a unique opportunity to evaluate whether seasonal differences in inversion frequencies that we observed in our data could be explained by sampling variance. We simulated a sampling scheme that emulated the empirical data to determine how often we observed seasonal differences in some inversion *I* as large as we observed empirically. To accomplish this, we resampled inversion-associated SNP frequencies based on the empirical coverage in each pool. For each pool, we assumed the true inversion frequency, *f*_*inv*_, is the midpoint of the seasonal point estimates. Using this assumed inversion frequency, we sampled the number of inversion-bearing chromosomes in a simulated pool as *chr*_*inv*_∼*binom* (*n*_*eff*_, *f*_*inv*_) where *n*_*e*_ is the effective pool size (as described above) of the simulated pool. For each inversion-associated SNP *j* in the empirical dataset, we simulated inversion-bearing reads based on the empirical coverage in the pool. Thus, 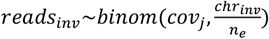 where *cov*_*j*_ is the coverage observed empirically at the *j*^*th*^ SNP. Just as we had in the empirical data, we then had a vector of simulated inversion-associated SNP frequencies. The mean of this vector was the simulated estimated inversion frequency of the resampled pool for inversion *i*, and the resampled seasonal estimate was calculated by weighting each resampled pool by effective ploidy. The P values shown in Table S5 are defined as the proportion of resampled seasonal differences that were greater than what we observed empirically. Essentially, we asked whether seasonal differences in inversion frequencies could be explained by sampling variance. We also used this sampling scheme to determine whether sampling variance could explain differences in inversion estimates by sampling location. Here, we noted the largest estimated inversion frequency difference and asked how often this difference was larger than what was observed empirically.

### Effective Population Size Estimate

In order to draw conclusions about frequency changes over the ∼35 year time period, we first needed to determine how much frequency change could be expected due to genetic drift. This expectation depends on the local size of the Rhode Island population, where a smaller population size would lead to a higher variance in the temporal frequency shift. To estimate this local population size, we simulated allele frequency trajectories of SNPs based on a simple Wright-Fisher model and fit the distribution of observed frequency changes to distributions of differing population sizes. The Wright-Fisher simulation emulated the empirical observations.

The site frequency spectrum of the old samples matched the simulated site frequency spectrum at the first time point. We then simulated 465 generations between the temporal samplings, corresponding to 15 generations per year (Turelli and Hoffmann 1995; Pool 2015) for 31 years. The SNP frequency at each generation, 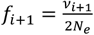, was drawn from a binomial distribution where *v*_*i+1*_∼*binom*(2*N*_*e*_, *f*_*i*_) i+1 is the number of individuals in the next generation bearing the allele, *N*_*e*_ is the effective population size in the simulation, and *f*_*i*_ is the allele frequency in generation *i*.

The sampling at the latter two time points emulated the sampling observed in the real data. For each SNP, the number of chromosomes in pool *j* that bear the allele (ψ) is drawn from a binomial distribution. Here, 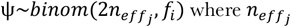 where 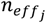 is the effective pool size of pool *j*. To further emulate the real data, coverage for all simulated pools is drawn from the empirical distribution of coverage. We then sampled reads bearing the allele, *ϕ*, based the coverage and 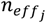 such that 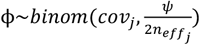 Simulated SNP frequency in pool *j* is then calculated as 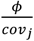 and the overall seasonal frequency is weighted by ploidy.

### Identification of Candidate Regions for Recent Directional Selection

The PBS statistic was used to quantify genetic differentiation specific to the newly collected seasonal samples when compared against the old population samples. Because differences in inversion frequencies between the sampling points can bias estimates of allele frequency differences, we weighted the old samples to match inversion frequencies of the pooled samples and corrected for sample size using Kish’s effective sample size (Kish 1965). PBS was applied in diversity-scaled genomic windows containing 200 nonsingleton SNPs in the Zambia sample of the *Drosophila* Genome Nexus. Simulations were run for autosomal and X chromosome data based on the model from Sprengelmeyer *et al*. (2020) and adding a North American arm with admixture (both proportion and timing) based on empirical data. This North American arm was admixed from the France and Cameroon arms of the model. We ran 2.5 million ms simulations (Hudson 2002) for each of 5 recombination bins corresponding to 0.5-1, 1-1.5, 1.5-2, 2-3, and greater than 3 cM/Mb. Regions of less than 0.5 cM/Mb were not included in our analysis due to limited resolution in localizing signals of elevated differentiation. For each simulation, we randomly chose a locus length from the distribution of empirical windows.

Recombination rate for each window was based on estimates from Comeron *et al*. (2012). The mutation rate for the autosomal model was 5.21e-9 and the X chromosome model was 5.07e-9 (Huang *et al*. 2016). Number of sampled chromosomes was also based on empirical data. To emulate pooled sequencing, we resampled simulated chromosomes based on coverage in the empirical data. With these simulations, we were able to assign a genome-wide P value for each window PBS by asking how many times simulated PBS values were greater than the observed PBS value. We assigned significance to any P value below the Bonferroni corrected critical value of 0.05/(number of genome-wide windows).

### Genome-wide Enrichment of PBS Outliers

In light of observed enrichments of window PBS and max-SNP PBS, we asked how many regions could be removed before the enrichment disappeared. We defined a region starting at a window with a low P value for window or max-SNP PBS and extending in each direction until hitting a string of 10 consecutive windows with a P value greater than 0.1. Windows were removed until the bin containing P values between 0 and 0.05 held no more windows than the bin containing P values between 0.05 and 0.1. For max-SNP PBS, because some SNP frequency estimates could be affected by differences in data types, we conservatively required the focal window of a region to also be an outlier (top 5%) in our “pseudo-pool” PBS scan (as described above) in order to be counted toward the number of removed regions. To ensure accuracy and minimize unaccounted for bias, we utilized two different approaches to removing regions, and they yielded similar results. In a deterministic approach, we iteratively removed the lowest P value region until no P value enrichment remained. In a random approach, we randomly chose a window whose P value was less than 0.05 and defined a region to remove around that window. This process was repeated 1,000 times and the average number of removed regions was noted.

### Gene Ontology Enrichment

The top 1% of PBS quantiles were considered outliers for GO enrichment analysis under the hypothesis that these outliers will be enriched for genuine targets of adaptation. GO enrichment was assessed as previously described in Pool *et al*. (2012). Two or more outlier windows were merged into the same outlier window region if they were separated by no more than four nonoutlier windows (to conservatively avoid counting the same selective sweep more than once). Locations of outlier regions were then randomly permuted, while maintaining their lengths, to properly account for the arrangement and lengths of genes in each functional category. Each outlier region was only allowed to count for a given GO category one time (from both the empirical and permuted outlier regions), to avoid spurious results from clusters of functionally linked paralogs. For each GO term, a raw P value was defined by the proportion of 1,000,000 randomized data sets in which a greater or equal number of outliers from that category was obtained. Then, by comparing across these randomized data sets, the lowest raw P value for each of them was obtained, and a threshold for analysis-wide significance was defined based on a minimum raw P value observed in 5% or fewer randomized data sets.

### Analysis of Frequency Change at the Cyp6a17/Cyp6a23 Locus

The *Cyp6a17*/*Cyp6a23* locus, a top resut from our genome-wide PBS scan, is known to have segregating structural variation involving a derived fusion of these paralogs into a single chimeric gene (Good et al. 2014). Identifying intact versus deleted alleles among the old isofemale strains was straightforward based on a roughly 20-fold difference in depth of coverage relative to genome-wide averages (Table S9). For the merged pooled data, we first calculated the ratio (*r*_*pool*_) of depth of coverage in the deletion region (*D*_*pool_focal*_) over the genome-wide average (*D*_*pool_genome*_), as well as the ratios of average depths seen in isofemale genomes inferred to have intact (*r*_*intact*_ = 1.2) or deleted (*r*_*deleted*_ = 0.0676) haplotypes. We then estimated the intact alelle frequency of the pooled data as (*r*_*pool*_ – *r*_*deleted*_) / (*r*_*intact*_ – *r*_*deleted*_), obtaining (0.646 - 0.0676)/(1.2 - 0.0676)=0.5108.

We used approximate Bayesian computation to estimate the strength of selection and the initial frequency of the intact *Cyp6a17* + *Cyp6a23* haplotype during the 1975 – 1983 sampling period (during which samples were taken during six years). We employed a very simple Wright-Fisher simulator and simulated 120 generations (15 generations over 8 years). For each simulation, we randomly selected a selection strength from a uniform distribution between 0 and 0.3 and an initial frequency from a uniform distribution between 0 and 0.2. We only accepted a simulation that exactly matched our empirical counts of an intact *Cyp6a17* gene. We stopped our simulation after 500,000 successes, which was around 100 billion simulations.

## Data Accessibility

All sequence data generated for this project is available from the NIH Short Read Archive under project SRX9688492, with specific sample numbers given in Table S1. All novel scripts used in this study have been uploaded to https://github.com/jeremy-lange/temporal.

## Acknowledgments

We are especially grateful to Rayla Temin for the gift of the original population sample of Providence lines. We also thank David Rand for the use of his fly room. This work benefitted substantially from computational resources and support from the UW-Madison Center for High Throughput Computing (CHTC). It was funded by USDA Hatch grant WIS01900 and by NIH grants R35 GM136306 and T32 GM007133.

