## Supplemental Figures for "A Population Genomic Assessment of Three Decades of Evolution in a Natural *Drosophila* Population"

**Figure S1:** Analysis of within-population Principle Component (PC) variability. Shown are the population-specific standard deviations for PC1 (A) and PC2 (B), along with the proportions of individual outliers for PC1 (C) and PC2 (D). Outliers were defined as PC values more than two population standard deviations from the population mean, and this analysis was limited to samples with sufficient numbers of individuals to estimate such a proportion with fair accuracy.

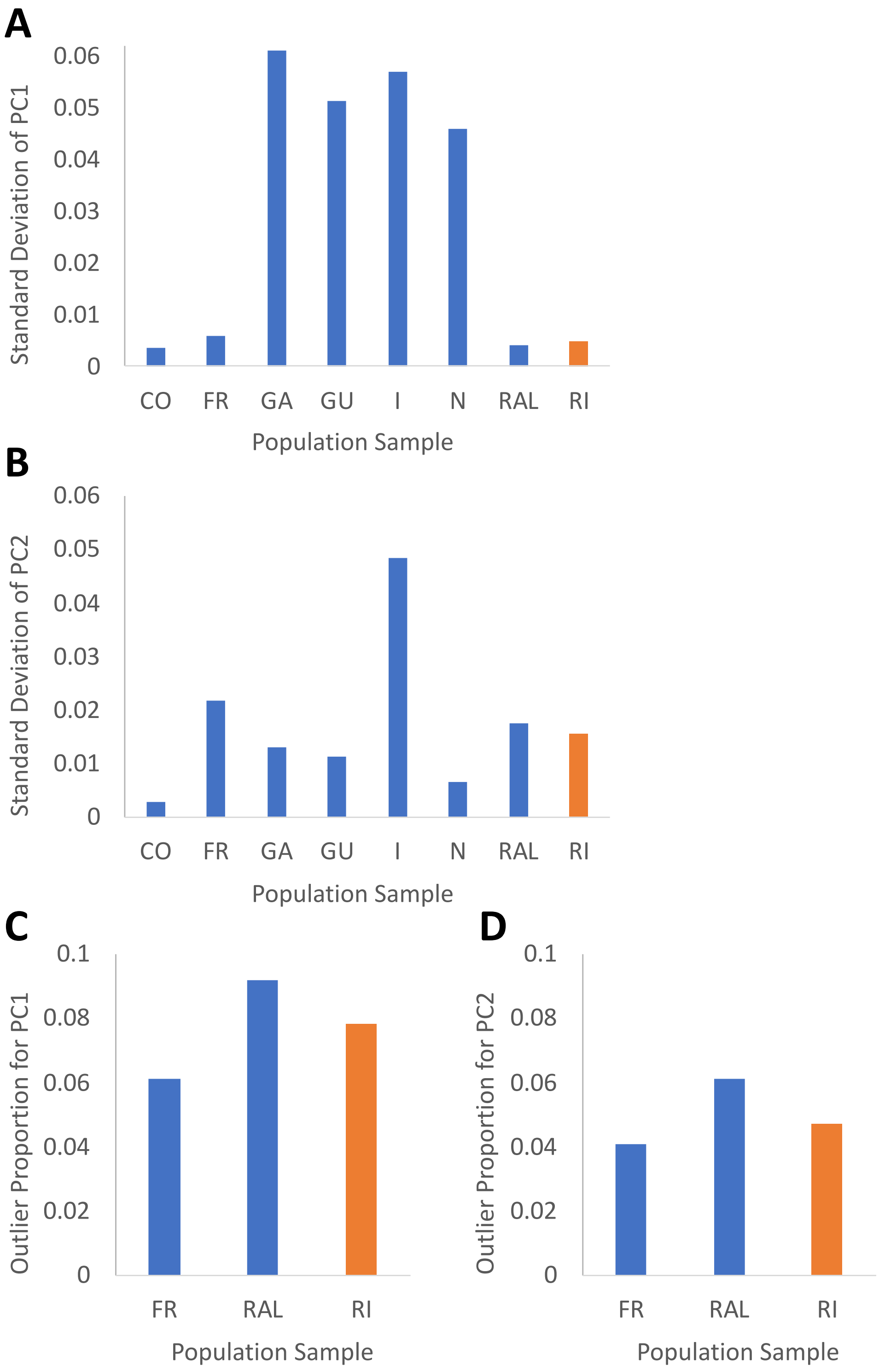

**Figure S2:** SNP PBS plots are shown for selected window PBS outliers that include the following genes: (A) *Hexo2*, (B) *Doa*, and (C) *CG4991* and *CG16700*.

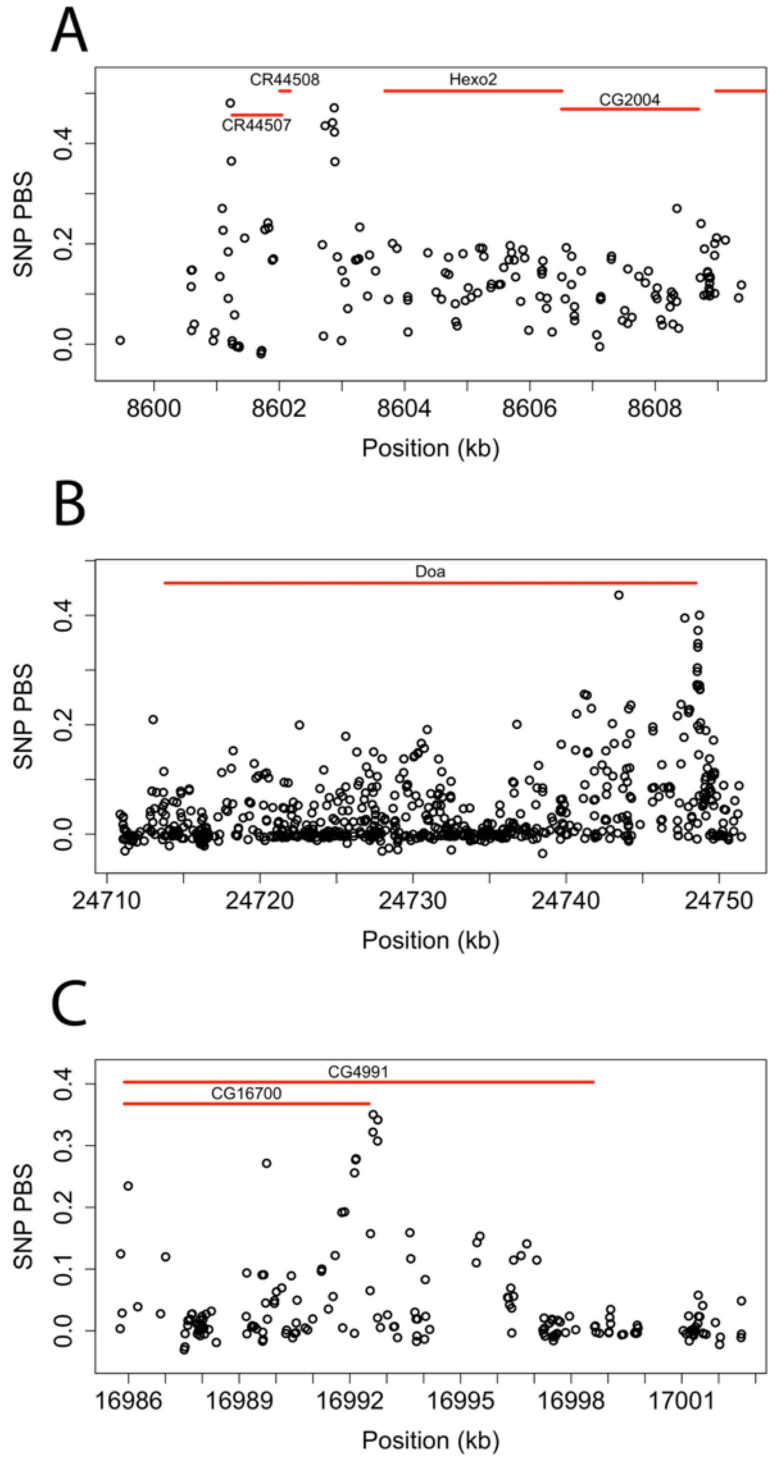

**Figure S3**: SNP PBS plots for a selection of regions with an outlier SNP but the window was not an outlier.

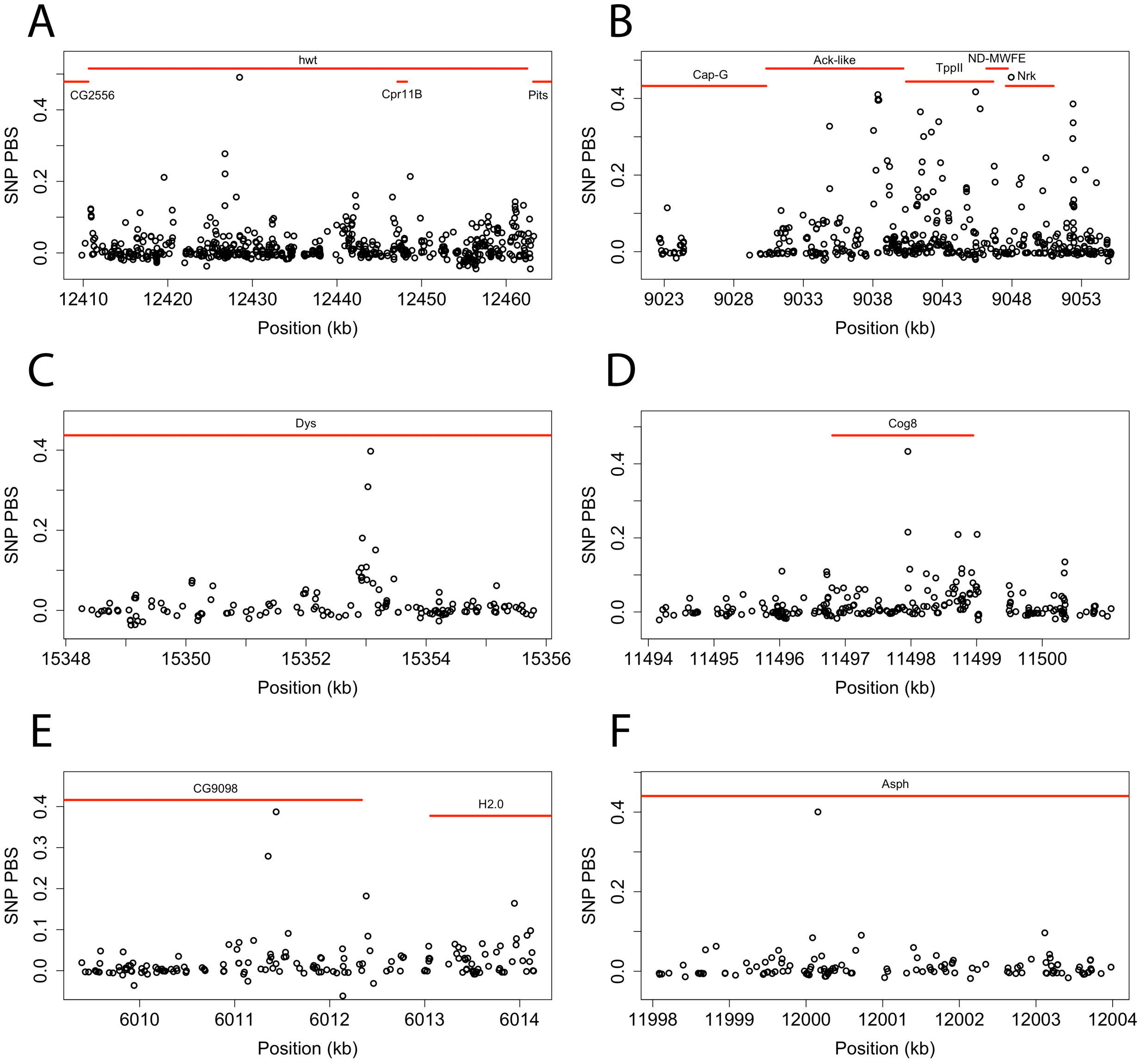
